# Stochastic variational variable selection for high-dimensional microbiome data

**DOI:** 10.1101/2021.10.04.462986

**Authors:** Tung Dang, Kie Kumaishi, Erika Usui, Shungo Kobori, Takumi Sato, Yusuke Toda, Yuji Yamasaki, Hisashi Tsujimoto, Yasunori Ichihashi, Hiroyoshi Iwata

## Abstract

**Background:** The rapid and accurate identification of a minimal-size core set of representative microbial species plays an important role in the clustering of microbial community data and interpretation of clustering results. However, the huge dimensionality of microbial metagenomics datasets is a major challenge for the existing methods such as Dirichlet multinomial mixture (DMM) models. In the framework of the existing methods, the computational burden of identifying a small number of representative species from a large number of observed species remains a challenge.

**Results:** We proposed a novel framework to improve the performance of the widely used DMM approach by combining three ideas: (i) we extended the finite DMM model to an infinite case by considering Dirichlet process mixtures and estimating the number of clusters as a random variables; (ii) we proposed an indicator variable to identify representative operational taxonomic units that substantially contribute to the differentiation among clusters; and (iii) to address the computational burden of high-dimensional microbiome data, we proposed a stochastic variational inference, which approximates the posterior distribution using a controllable distribution called variational distribution, and stochastic optimization algorithms for fast computation. Using the proposed method, stochastic variational variable selection (SVVS), we analyzed the root microbiome data collected in our soybean field experiment, the human gut microbiome data from three published datasets of large-scale case-control studies and the healthy human microbiome data from the Human Microbiome Project.

**Conclusions:** SVVS demonstrated a better performance and significantly faster computation than those of the existing methods in all cases of testing datasets. In particular, SVVS is the only method that can analyze massive high-dimensional microbial data with more than 50,000 microbial species and 1,000 samples. Furthermore, recent microbiome studies have suggested that selection of the microbial species used as a core set is important.

## Background

The development of metagenomic high-throughput sequencing technologies has provided a rapid and sensitive method for the discovery of human and soil microbial communities. Accordingly, our understanding of the impact of the gut microbiota on the human body [1, 2] and the significance of bacterial ecology in the global biogeochemical nutrient cycle [3] has greatly expanded. There are two major types of microbial metagenomic data: 16S ribosomal RNA genes and shotgun metagenomics. In this study, we focused on the analysis of the 16S ribosomal RNA gene as an example, although shotgun metagenomics data can be analyzed in a relatively similar manner. One standard approach is to transform the 16S rRNA gene of the bacteria in the samples into operational taxonomic units (OTUs) using a preprocessing method, such as QIIME [4]. Next, using the transformed data sets, we aimed to identify the differences in the microbial composition among multiple groups of communities and to elucidate the relationship between these groups.

Considering the heterogeneous pattern of sample-to-sample variability in the microbiome data, various model-based approaches have been proposed for clustering microbiome samples. The finite Dirichlet multinomial mixture (DMM) model is one of the most widely used methods [5]. The main ideas behind the DMM model are as follows. First, a multinomial sampling scheme is adopted for the OTU count data, and then a mixture of Dirichlet components is considered as the natural prior for the parameters of the multinomial distribution. This approach helps avoid the disadvantages of previous methods, assuming that all samples in a cohort are generated from a single community profile, and allows each community to be considered a mixture of multiple communities, which can be described by a vector generated by one of the finite Dirichlet mixture components with different hyperparameters. Therefore, the flexibility of the DMM model with respect to model dimensionality makes it well suited for capturing many different sub-community structures. Such methods are of great practical importance and have been used to assess the potential associations of the microbiome community in studies on human health and disease [6, 7], microbiome genome-wide association [8], and animal microbiomes [9].

The first step in an analysis with a conventional DMM model is to determine the number of microbiome clusters, that is, the metacommunities biologically required to explain the observations. A fully Bayesian model selection through Laplace approximation [5] has been proposed to consider all possible values for the number of metacommunities up to a certain maximum value. However, the optimization of this number via this approach is computationally prohibitive and may cause poor performance when the dimensionality of the microbiome datasets is high. Moreover, it is worth mentioning that in the DMM model, all OTUs are assumed to be equally significant in the clustering analysis. However, this is not realistic in practical analysis because of the thousands or tens of thousands of OTUs; a large number might be irrelevant and might not significantly contribute to the identification (or characterization) of the microbiome clusters.

Recently, various potential approaches have been proposed to estimate the parameters of a nonparametric Bayesian unsupervised variable selection in the field of computer science, such as a hybrid framework based on a combination of Gibbs sampling and Metropolis-Hastings algorithms that appropriately accounts for the conditional independence relationships between latent variables and model parameters [10, 11]. An important advantage of Markov chain Monte Carlo (MCMC) solutions is the yielding of a robust and stable estimation by including the observed data in the computational processes. However, the computational burden of MCMC solutions is prohibitive for inference given the large dimensionality of microbial metagenomics datasets, and it can be very difficult to diagnose their convergence.

In this study, we proposed a novel approach that overcomes the challenges described above and achieves feasible computational ability for a personal computer. The main contributions of this study are threefold. First, we extended the finite DMM model by proposing a Bayesian nonparametric approach based on a countable infinite mixture model coupled with variable selection. In our approach, the number of clusters (metacommunities) is not fixed a priori and is itself a free parameter of inference under the truncated stick-breaking representation of the Dirichlet process prior to the space of mixture metacommunities [12, 13, 14]. This solution can overcome the difficulty of choosing an appropriate number of clusters based on the data. Moreover, our method enabled the estimation of the significant contributions of OTUs to detect a minimum core set of OTUs that characterize clusters and maximize the identification ability of the clusters. Second, to overcome the current computational difficulties related to deterministic learning and MCMC approaches, we proposed a stochastic variational inference (SVI) method [15, 16, 17], which was originally used in statistical physics to approximate intractable integrals and has been successfully used in a wide variety of applications for analyzing large datasets related to population genetics [18, 19] and phylogenetics [20, 21, 22]. It is worth noting that an analytically tractable solution for calculating the posterior distributions has been provided to avoid expensive computations of the numerical approximations in MCMC approaches [23, 24].

Finally, to test the performance of the proposed approach, we used two types of 16S rRNA gene amplicon sequencing microbiome data. The first type included several datasets in which numerous groups were known and the samples were clearly labeled. Thus, we could measure the similarity between the truth clusters and inferred clusters to compare the accuracies of the different approaches. In this study, we analyzed the environmental microbiome data of soybean genetic resources collected in our field experiments. The dataset included 196 and 197 rhizosphere samples, which contained 888 OTUs from a drought irrigation and control conditions, respectively. Moreover, our proposed approach was applied to three published casecontrol 16S rRNA gene amplicon sequencing datasets of the human gut microbiome [25, 26, 27]. Specifically, the first database included 3,347 OTUs for *Clostridium difficile* infection (CDI) from 338 individuals, including 89 individuals infected with CDI (cases), 89 individuals with diarrhea who tested negative for CDI (diarrhea controls), and 155 non-diarrheal controls [25]. Moreover, inflammatory bowel disease (IBD) data and obesity (OB) data provide numerous OTUs (approximately 10,000 and 50,000, respectively), which challenge the computational capability of stochastic variational variable selection (SVVS) [26, 27]. Conversely, various studies of the healthy human microbiome have shown that if the number of groups of samples is still unknown, the identification of the clusters (referred to as enterotypes) is difficult [28]. Therefore, in the second type, we analyzed a dataset in which the number of groups was unknown. This dataset included the stool samples from the Human Microbiome Project (HMP), specifically the *HMP16SData* package, which had 319 samples and 11,747 OTUs [29], to identify the number of distinct clusters (or enterotypes).

## Materials and Methods

### The finite Dirichlet multinomial mixture model

First, the finite DMM model that describes the heterogeneity of cross-sample variability among microbiome species was briefly reviewed [5]. This model allows a dataset to be generated by a mixture of K metacommunities instead of a single metacommunity. The key concepts behind the DMM model are as follows:

Given a microbiome dataset consisting of *N* community samples and *S* OTUs (or species), the observed count of the *i^th^* community for *j^th^* OTU was denoted as *X_ij_*(*i* = 1,…, *N*; *j* = 1, …, *S*). The total number of counts (i.e., sequence reads) from the *i^th^* community sample was 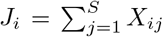. The DMM model [5] considered a vector 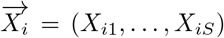, drawn from a multinomial distribution with community vector 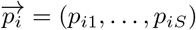 as follows:

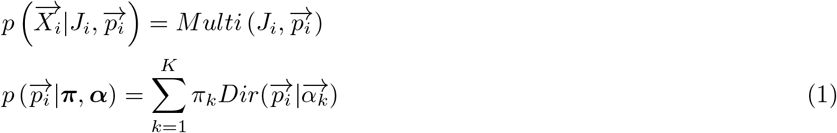

where *p_ij_* is the probability that a single read in the *i^th^* community belongs to the *j^th^* OTU; 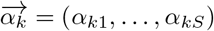 are the parameters of the Dirichlet distribution representing the *k^th^* metacommunity (or cluster); and ***π*** = (*π*_1_,…, *π_K_*) represents the mixing coefficients with 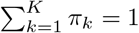, *π_k_* ≥ 0 for *k* ∈ (1,…, *K*). It is worth noting that the finite DMM model examined a case where the number of metacommunities, K, was fixed. Each sample was assumed to be drawn from each unique community vector 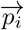, which was derived from one of the K metacommunities. Therefore, Equation (1) can be rewritten by marginalizing the multinomial parameters as follows [5]:

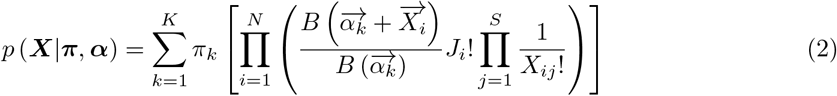

where the function B is the multinomial beta function 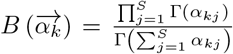 and 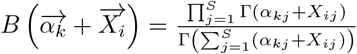

### The infinite Dirichlet multinomial mixture model with variable selection

The goal was to consider the number of metacommunities (*K*) as a random variable. To achieve this, it was assumed that the prior distribution of the mixing coefficients ***π*** followed a Dirichlet process prior [12]. The stick-breaking representation [13] [14], which is a straightforward constructive definition of the Dirichlet process, was adopted to construct the infinite DMM model proposed in this study. This was defined as follows:

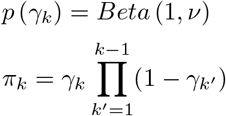

where *π_k_* is the mixing proportion of an infinite number of successively broken sticks, and the unit length of sticks that correspond to the *k^th^* metacommunity (or cluster), with (*k* ∈ [1,…, *K*]), and *ν* represents the total mass parameter of the Dirichlet process. It was assumed that each community sample 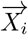 was generated from the DMM model with a countably infinite of number of clusters (or metacommunities). Therefore, the Equation (2) can be rewritten as

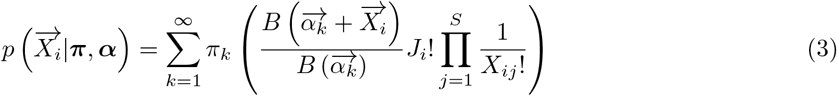

Next, we introduced the allocation variable 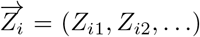, such *Z_ik_* ∈ [0, 1] and *Z_ik_* = 1 if 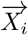 belongs to the metacommunity *k* (i.e., the cluster *k*) or 0. Moreover, all OTUs in the DMM model were assumed to be equally important for clustering microbial community data. However, this is not realistic in microbiome studies, because numerous microbiome species (which can be reflected in OTUs) and functions might be irrelevant and significantly influence the performance of clustering algorithms [30]. To overcome this problem, we proposed that the count of a given OTU, *X_ij_*, be generated from a mixture of two Dirichlet-multinomial distributions; the first one was assumed to generate a core set of the most significant microbial OTUs and was different for each single metacommunity, and the second one was assumed to generate the unimportant OTUs and was common to all multiple metacommunities. Thus, we could write the likelihood of the observed microbiome dataset ***X*** following the infinite DMM model with microbiome OTU selection as follows:

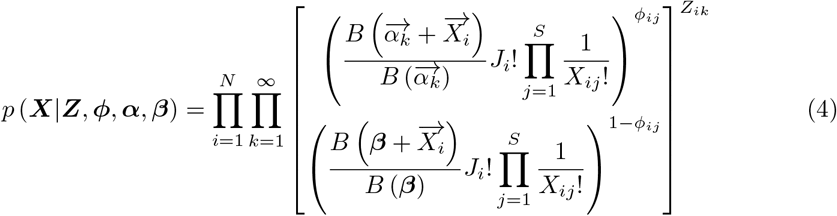

where *ϕ_ij_* is an indicator variable, such that *ϕ_ij_* = 1 indicates that the *j^th^* OTU of the *i^th^* community is important in the *k^th^* cluster and follows a Dirichlet multinomial distribution with ***α***, and *ϕ_ij_* = 0 denotes that the *j^th^* OTU of *i^th^* the community is unimportant in the *k^th^* cluster and follows a Dirichlet multinomial distribution with ***β***. 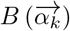 and 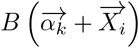 are the multinomial beta functions for a core set of OTUs that significantly represent the cluster. For unimportant species, the multinomial beta functions were 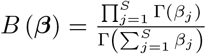 and 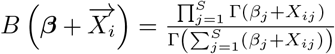. The prior distribution of the indicator variable of microbiome selection ***ϕ*** was defined as follows:

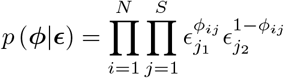

where each *ϕ_ij_* follows a Bernoulli distribution such that *p* (*ϕ_ij_* = 1) = *ϵ*_*j*1_ and *p*(*ϕ_ij_* = 0) = *ϵ*_*j*2_ with *ϵ*_*j*1_ + *ϵ*_*j*2_ = 1 [11]. Furthermore, the Dirichlet distributions are considered for (*ϵ*_*j*1_, *ϵ*_*j*2_) with the hyperparameters (*ξ*_1_, *ξ*_2_) [31]. The prior distributions of ***α*** and ***β*** follow the Dirichlet distributions with hyperparameters ***ζ*** and ***η***.

### Stochastic variational variable selection approach

In this study, we proposed an SVI method [15] [16] [17] for the infinite DMM model with the selection of variables (i.e., the counts of OTUs). The basic idea of variational learning in the Bayesian framework is to approximate the posterior distribution using a computationally tractable function called the variational distribution. The variational parameter, which specifies the variational distribution, is estimated by minimizing the Kullback-Leibler (KL) divergence of the posterior distribution to the variational distribution. As a result, the posterior distribution is estimated by numerical optimization with the triggering of the simulation approaches, such as MCMC algorithms.

Given the observed count dataset ***X***, the infinite DMM model has a set of parameters (Ξ), which consists of the unit length of the stick (**γ**), the allocation variable (***Z***) of the prior Dirichlet, the indicator variable of the OTU selection (***ϕ***), and the Dirichlet parameters (***α, β***). Variational inference approximates the true intractable posterior distribution *p*(Ξ|***X***) by an element of a tractable family of probability distributions *q*(Ξ|Θ) called the variational distribution. This distribution is parameterized by free parameters, called variational parameters Θ. Variational inference estimates these parameters to find a distribution close to the true intractable posterior distribution of interest. The distance between the distributions *p*(Ξ|***X***) and *q*(Ξ|Θ) is evaluated using KL divergence, represented as follows:

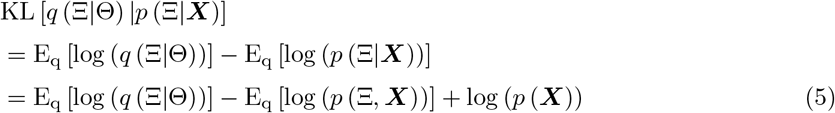

The log marginal probability log (*p*(***X***)) in Equation (5), which causes computational difficulty in the Bayesian framework, can be treated as a constant term in the numerical optimization for estimating the variational parameters as follows:

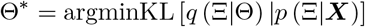

In addition, the term log (*p*(***X***)) can be decomposed as 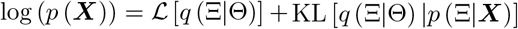. The variational inference maximizes the computationally feasible target function as follows:

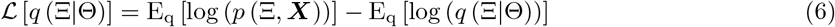

where Equation (6) is the Evidence Lower Bound (ELBO) [15]. 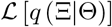 can be considered a lower bound for log (*p*(***X***)). Perceptibly, the maximization of ELBO equals the minimization of KL divergence, that is, when the variational distribution *q*(Ξ|Θ) approximates the true posterior distribution *p*(Ξ|***X***). However, the fact indices of the computation of the true posterior distribution are intractable, and direct application of the variational framework is infeasible. Therefore, a mean-field framework was adopted in order to factorize the posterior distribution into disjoint tractable distributions. According to the factorization assumption of mean-field variational approximations [16] [17], each variable in the variational distribution *q*(Ξ|Θ) is independent. Furthermore, we considered truncated stick-breaking representations for the DMM model at the largest possible number of metacommunities (or clusters) K_max_. It is worth mentioning that the variational framework can optimize the value of truncation K_max_ because it becomes a variational parameter. The family of variational distributions in the infinite DMM model with the selection of representative OTUs can be expressed as follows:

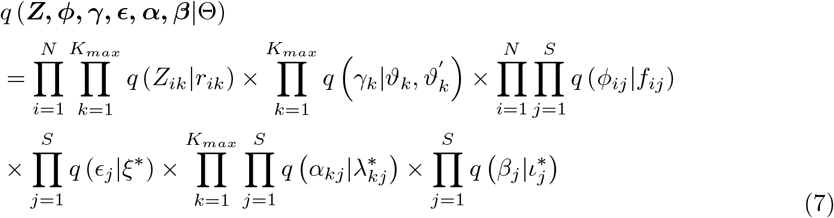

where

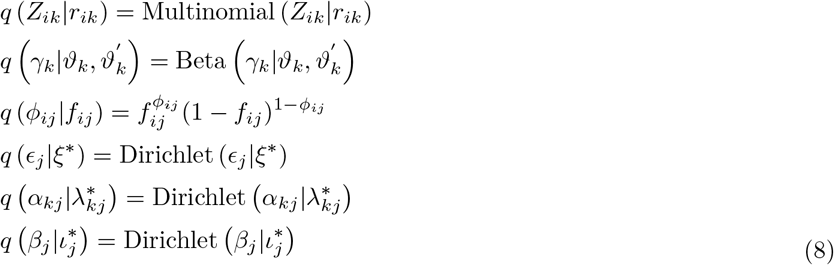

The set of free variational parameters Θ includes 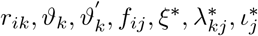. We considered the variational distributions from exponential families to guarantee tractable computations of expectations.

We adopted (SVI) in order to estimate each variational parameter in our proposed method [17] [20]. The key idea of this inference is to divide the variational variables into two subgroups: the local variables [Ξ_*l*_ ∈ (***Z, ϕ***)], which are per-datapoint latent variables, and the global variables [Ξ_*g*_ ∈ (***γ, ϵ, α, β***)], which potentially control all the data. The *i^th^* local variable *Z_ik_* of the mixture component, which represents the allocation of sample i, is governed by the local variational parameter *r_ik_*. In addition, the local variational parameter *f_ij_* was proposed to capture the *i^th^* local variable *ϕ_ij_*, which represents the selection situation of the *j^th^* OTU in the *i^th^* community. The coordinate ascent algorithm was considered to overcome the optimization problems of these variational variables [16] [17]. The main idea of this approach was to optimize each factor of the mean-field variational distribution while fixing the others. Then, the logarithm of the optimal value of *q*(*Z_ik_*) is proportional to the expected logarithm of the joint distribution as follows:

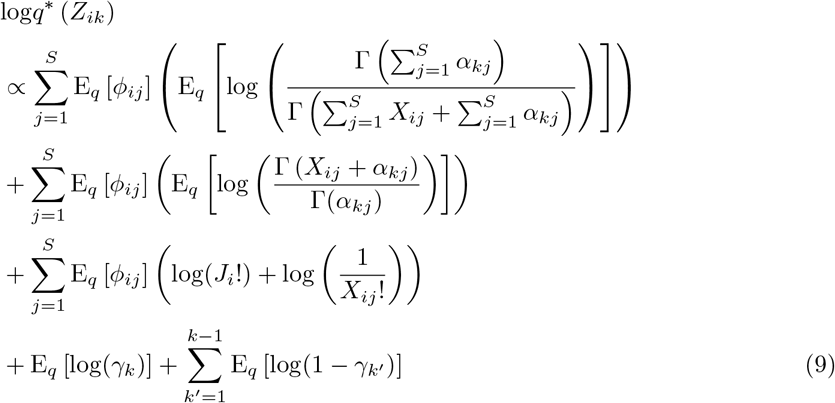

As *γ_k_* follows a beta distribution, we can obtain the analytically tractable solutions for E_*q*_[log(*γ_k_*)] and E_*q*_ [log(1 – *γ_k′_*)]. However, the first and second terms of Equation (9) do not have the same form as the logarithm of the Dirichlet prior distribution. Thus, analytically tractable solutions cannot be obtained directly. The intractable computation of expectations can be resolved using the Metropolis-Hastings algorithm and numerical integration. Nevertheless, the simulation approaches significantly increase the computational burden in the huge dimensionality of microbial metagenomics datasets [5]. Therefore, we adopted the Taylor expansion to obtain the nearly optimal analytically tractable solutions for the first and second terms of Equation (9), such that the computational burdens were avoided [32] [23] [24]. A nearly optimal analytically tractable value of q (*ϕ_ij_*) could be obtained using the proposed approach. The mathematical details of the Taylor expansion and variational objective functions are provided in the Supplementary Material.

The global variational parameters 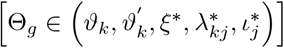 were proposed to govern the global variable Ξ_*g*_. The SVI approach uses the stochastic gradient ascent in order to estimate the global variational parameters [17]. Furthermore, the natural gradients have been adopted to account for the geometric structure of probability parameters [33] [34] [35]. It is worth noting that the natural gradients are not only cheaper computations but also have faster convergence than that of standard gradients. The SVI is based on the stochastic approximation approach to iteratively generate subsampled datasets that are used to update the values of the local and global variational parameters. The main advantage of these computational strategies is that they guarantee algorithms that avoid shallow local optima for complex objective functions.

For the infinite DMM model, we generated a uniform a dataset 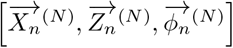 that was formed by N replicated from the microbiome community sample 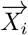, allocation variable 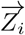, and indicator variable 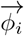 at each iteration. Next, noisy estimates of the natural gradient were computed with respect to each global variational parameter Θ_*g*_ given N replicates of the sampled data point. Using these gradients, the values of Θ_*g*_ were updated at each t iteration given the local variational parameters [Θ_*l*_ ∈ (*r_ik_, f_ij_*)] as follows:

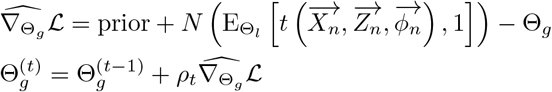

where t(.) denotes the sufficient statistics in the exponential family and *ρ_t_* denotes the step size at iteration t. Owing to the subsampling strategies, the SVI significantly accelerated the computational processes by avoiding expensive sums in the ELBO when the dimensionality of the microbial metagenomics was large. The mathematical explanations of the SVI are described in the Supplementary Material.

### Criteria to evaluate the performance of the approaches

We used the Adjusted Rand Index (ARI) [36] in order to measure the similarity between the truth (or known) clusters and clusters inferred by various algorithms. Given a dataset of ***X*** with n total samples, ***Z*** = [*Z*_1_,…, *Z_k_*] denotes the true cluster memberships of ***X*** into k clusters, and 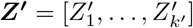 denotes an inferred cluster membership of ***X*** into k’ clusters. The Rand Index (RI) was calculated as follows:

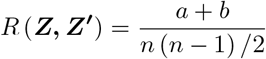

where a denotes the number of times a pair of samples was assigned to the same cluster in ***Z*** and ***Z***′, and b denotes the number of times a pair of samples was assigned to different clusters in ***Z*** and ***Z***′. The RI values were in the range of [0,1], where 1 represented a perfect similarity between the truth and inferred clusters. The ARI was proposed to normalize the difference between the RI and its expected value as follows:

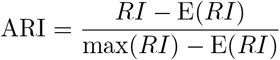

where E(*RI*) was the expected value of the RI.

### Database description

#### Tottori, Japan study

##### Field experiment

A total of 198 soybean accessions registered in the National Agriculture and Food Research Organization Genebank (https://www.gene.affrc.go.jp/) were used. The field trial was conducted in 2018 in an experimental field with sandy soil at the Arid Land Research Center, Tottori University (35°32’ N, 134°12’ E, 14 m above sea level). Each plot consisted of four plants, and the distances between two rows, two plots, and two individuals were 50, 80, and 20 cm, respectively. Sowing was performed at the beginning of July, followed by thinning after two weeks. Fertilizer (15, 6.0, 20, 11, and 7.0 g m-2 of N, P, K, Mg, and Ca, respectively) was applied to the field before sowing. White mulching sheets (Dupont, Wilmington, DE, USA) were laid to prevent rainwater infiltration and to control soil conditions with artificial irrigation. Watering tubes were installed under the sheets to irrigate the fields. Two watering treatments, non-watered treatment and well-watered treatments, were used to evaluate the influence of drought and control conditions. The watering treatment started after thinning every year, two weeks after sowing. Artificial irrigation was applied for 5 h daily (7:00–9:00, 12:00–14:00, and 16:00–17:00).

##### DNA extraction

DNA extraction was performed according to our custom protocol using magnetic beads [37]. Briefly, the collected root tissues were ground to a fine powder using a multi-beads shocker (MB2200(S); Yasui Kikai Co. Osaka, Japan). For each of the collected tissue samples, 500 mg of the powdered sample was transferred into a 1.5 mL tube cooled by liquid nitrogen. Then, 1 mL of lysate binding buffer (1 M LiCl, Cat. #L7026-500ML, Sigma-Aldrich, St. Louis, MO, USA; 100 mM Tris-HCl, Cat. #318-90225, Wako Pure Chemical Corporation, OSA, Japan; 1% SDS, Cat. #313-90275, Wako Pure Chemical Corporation; 10 mM EDTA pH 8.0, Cat. #311-90075, Wako Pure Chemical Corporation; Antifoam A, Cat. #A5633-25G, Sigma-Aldrich; 5 mM DTT, Cat. #048-29224, Wako Pure Chemical Corporation; 11.2 M 3-Mercapto-1,2-propanediol, Cat. #139-16452, Wako Pure Chemical Corporation; and DNase/RNase-free H2O, Cat. #10977015, Thermo Fisher Scientific, Waltham, MA, USA) [38] was added to the sample, which was then homogenized by vortexing, followed by incubation at room temperature (22°C) for 5 min. The tube was centrifuged at 15,000 rpm for 10 min at room temperature (22°C), and the supernatant was transferred to a new 1.5 mL tube. Subsequently, 50 *μ*L of LBB lysate was added to 1.5 mL tubes, and an equal amount of AMPure XP beads was added, followed by incubation at room temperature (22°C) for 5 min after vortexing. The mixture was placed on a magnetic station for 5 min and the supernatant was removed. The magnetic beads were washed twice with 200 *μ*L of 80% ethanol. Finally, DNA was eluted with 20 *μ*L of 10 mM Tris-HCl (pH 7.5).

##### 16S rRNA gene amplicon sequencing

Library preparation using a two-step PCR amplification protocol has been reported in our previous publication [37, 38]. Briefly, the V4 region of the bacterial 16S rRNA gene was amplified using 515f and 806rB primers (forward primer: 5’-TCG TCG GCA GCG TCA GAT GTG TAT AAG AGA CAG- [3–6-mer Ns] – GTG YCA GCM GCC GCG GTA A −3’; reverse primer: 5’-GTC TCG TGG GCT CGG AGA TGT GTA TAA GAG ACA G [3–6-mer Ns] - GGA CTA CNV GGG TWT CTA AT −3’) [39, 40]. Each sample (1 *μ*L of 10-fold diluted DNA) was amplified in a 10 *μ*L reaction volume containing 0.2 U KOD FX Neo DNA polymerase (TOYOBO Co., Ltd., Osaka, Japan), 2 × PCR buffer (TOYOBO Co., Ltd.), 0.4 mM dNTPs (TOYOBO Co., Ltd.), 0.2 *μ*M forward and reverse primers, and 1 *μ*M blocking primers (mPNA and pPNA; PNA BIO, Inc., Newbury Park, CA, USA). PCR was performed using the following conditions: 94°C for 2 min, followed by 35 cycles at 98°C for 10 s, 78°C for 10 s, 55°C for 30 s, 68°C for 50 s, and a final extension at 68°C for 5 min (ramp rate = 1°C/s).

The first PCR products were purified using a mixture of exonuclease and alkaline phosphatase. Two *μ*L of ExoSAP-IT Express (Cat #75001.1.EA; Thermo Fisher Scientific) was added to 5 *μ*L of the product obtained from the first PCR, and the mixture was incubated at 37°C for 4 min, followed by incubation at 80°C for 1 min.

The second PCR was conducted using the following primers: forward primer: 5’-AAT GAT ACG GCG ACC ACC GAG ATC TAC AC - [8-mer index] - TCG TCG GCA GCG TC −3’, and reverse primer: 5’- CAA GCA GAA GAC GGC ATA CGA GAT - [8-mer index] - GTC TCG TGG GCT CGG −3’ (Toju and Baba, 2018). Each sample (0.8 *μ*L of purified product from the first PCR) was amplified in a 10 *μ*L reaction volume containing 0.2 U KOD FX Neo DNA polymerase (TOYOBO Co., Ltd.), 2 × PCR buffer (TOYOBO Co., Ltd.), 0.4 mM dNTPs (TOYOBO Co., Ltd.), 0.3 *μ*M forward and reverse primers, and 1 *μ*M blocking primers (mPNA and pPNA). PCR was performed as follows: 94°C for 2 min, followed by eight cycles at 98°C for 10 s, 78°C for 10 s, 55°C for 30 s, 68°C for 50 s, and a final extension at 68°C for 5 min (ramp rate = 1°C/s). Following amplification, the PCR products for each sample were cleaned and size-selected using AMPure XP beads and washed twice with 80% ethanol. The libraries were eluted from the pellet with 10 *μ*L of 10 mM Tris-HCl pH 7.5, quantified using a microplate photometer (Infinite 200 PRO M Nano+, TECAN Japan Co., Ltd., Kanagawa, Japan), and pooled into a single library in equal molar quantities. The pooled library was sequenced on an Illumina MiSeq platform using a 2× 300-bp MiSeq Reagent Nano Kit v3 (Illumina, San Diego, CA, USA).

##### Taxonomic profiling and data preprocessing

Sequencing primers and short (<150 bp) reads were removed using Cutadapt v.1.18. The primer-free FASTQ files were further processed using R v3.6.0 with DADA2 v.1.14.0 [41]. First, the FASTQ file was trimmed, with a truncation length of 240 for forward reads and 160 for reverse reads. These reads underwent further quality filtering, as the error rates were calculated and removed from the duplicated reads. Forward and reverse sequences were merged. Next, potentially chimeric sequences and sequencing errors were removed. Taxonomy was assigned using the SILVA v.132 training set [42]. The filtered matrix was rarefied to 3,000 reads per sample using the vegan 2.5-6 package [43] and then used to calculate Bray-Curtis distance. We denoted Data set A as our Tottori data set.

#### Study inclusion and data acquisition

We also employed case–control 16S amplicon sequencing from three published human microbiome datasets spanning three different disease states: *Clostridium difficile* infection (CDI) [25], inflammatory bowel disease (IBD) [27], and obesity (OB) [26]. These datasets are available in the MicrobiomeHD database [44]. Dataset B represented the CDI dataset, which included 183 diarrheal stool samples from the 94 individuals with CDI, 89 diarrheal control samples, 155 non-diarrheal control stool samples, and 3,347 microbiome species (or OTUs). Dataset C represented the IBD dataset, which included 146 IBD case samples, 16 non-IBD control samples and 10,119 microbiome species (or OTUs). Dataset D represented the OB dataset, which was the largest and most challenging. There were 1,081 fecal samples from 977 individuals and 55,964 microbiome species (or OTUs). Finally, we used the variable region 3-5 (V35) of the 16S rRNA gene sequence dataset from the *HMP16SData* package to study the considerable variation in the composition of the healthy human microbiome [29]. Dataset E represented the data of stool community types from the *HMP16SData* package, which included 319 samples and 11,747 microbiome species (or OTUs).

### Open-source software

The software was implemented in Python and used standard libraries, such as NumPy and SciPy, for mathematical computations. The software inputs microbiome count data in a CSV file and outputs the inferred clusters and a core set of selected OTUs. The main options in the software tool are the maximum number of clusters, which pose limitations in estimating the number of clusters, and the number of OTUs that users want to select. We kept the convergence criterion fixed across all datasets in this study. The SVVS algorithm stopped when the change in the ELBO computations was less than 1e-3. The number of iterations was modified for datasets notably smaller or larger in scale than those considered in this study. This is a tunable option in the software. The software is available at https://github.com/tungtokyo1108/SVVS

## Results

### Runtime performance and physical memory of the computational system

An important advantage of SVVS over conventional DMM approaches is that the computational time and memory required for calculations can be greatly reduced. To evaluate the computational time and memory of the different approaches, we diversified the sample sizes and number of OTUs in the sample datasets. The scalability of the methods was specifically demonstrated in cases of datasets C and D; meanwhile, datasets A and B were selected to compare their accuracies. We followed the Laplace approximation to the model evidence and default values of the DirichletMultinomial 1.34.0 package in R to determine the number of clusters K for the finite DMM model [5] [45]. Our proposed method does not require selection of the number of clusters because the number of clusters is estimated as a random variable. Our Python implementations of SVVS for the infinite DMM model were used to analyze all real datasets. Tables 1 and 2 compare the computational time and physical memory required for the calculation between the SVVS algorithm of the infinite DMM model and the EM algorithm of the finite DMM model. We found that SVVS was able to considerably reduce run times and physical memories for datasets A, B, and C. SVVS was the only approach that was able to analyze dataset D, which is a large dataset of more than 50,000 OTUs and 1,000 samples. In addition, the time complexity and memory of each of the above methods were found to increase regularly with the number of OTUs and samples. SVVS used the iterative optimization algorithms to estimate the parameters; thus, a convergence criterion was used to implement a stopping rule (Supplementary Material).

**Table 1.** Running time of the two approaches on the real data sets. Note: All algorithms were run on a personal computer (Intel® Xeon® Gold 6230 Processor 2.10 GHz × 2, 40 cores, 2 threads per core, 128 Gb RAM) under Ubuntu 20.04.1 LTS.

| Datasets | Finite DMM with EM algorithm | Infinite DMM with SVVS algorithm |
| --- | --- | --- |
| A | 17.63 min | 2.68 min |
| B | 2.75 h | 13.25 min |
| C | 3.37 d | 30 min |
| D | Failed | 5 h |

**Table 2.** Physical memories of the two approaches on the real datasets

| Datasets | Finite DMM with EM algorithm | Infinite DMM with SVVS algorithm |
| --- | --- | --- |
| A | 1.02 Gbs | 0.186 Gbs |
| B | 3.5 Gbs | 1.65 Gbs |
| C | 15 Gbs | 4.5 Gbs |
| D | over 128 Gbs | 45 Gbs |

### The SVVS improves the accuracy of the approach

Table 3 compares the number of clusters predicted using the two approaches. Both the SVVS algorithm of the infinite DMM model and the EM algorithm of the finite DMM model obtained the correct numbers of clusters for datasets A and B. However, the number of OTUs was significantly larger in datasets C and D, and the SVVS approach achieved the most accurate predictions. Moreover, Table 4 compares the ARI values of the two methods. The SVVS algorithm of the infinite DMM model demonstrated a better performance than the conventional finite DMM model for all real datasets. Specifically, SVVS showed the highest ARI value (ARI = 0.98) for dataset A; coversely, the ARI value of the finite DMM with the EM algorithm was 0.76. For dataset B, the ARI values were slightly reduced in the performance of the SVVS (ARI = 0.66) and EM algorithms (ARI = 0.44). The number of OTUs in dataset B (3,347 OTUs) was significantly larger than that in dataset A (888). The small sample size and large number of OTUs in dataset C considerably influenced the performance of the two approaches. The lowest ARI values were observed for the SVVS (ARI = 0.48) and EM algorithms (ARI = 0.21). However, the ARI value of the SVVS approach increased slightly in dataset D when the number of samples was increased. The EM algorithm of the finite DMM model could not complete its estimation in dataset D, in which the dimensionality of the microbial data was the highest. Furthermore, to address graphical visualizations for the cluster labels that were predicted by the SVVS approach for the dataset A, we used non-metric multidimensional scaling (NMDS), which was performed on the unweighted UniFrac distance, to generate two-dimensional positions for community samples. Unweighted UniFrac distance, and NMDS functions are available in the phyloseq package [46]. Figures 1a and b show that the two groups of dataset A are separated by the two approaches.

**Figure 1.**
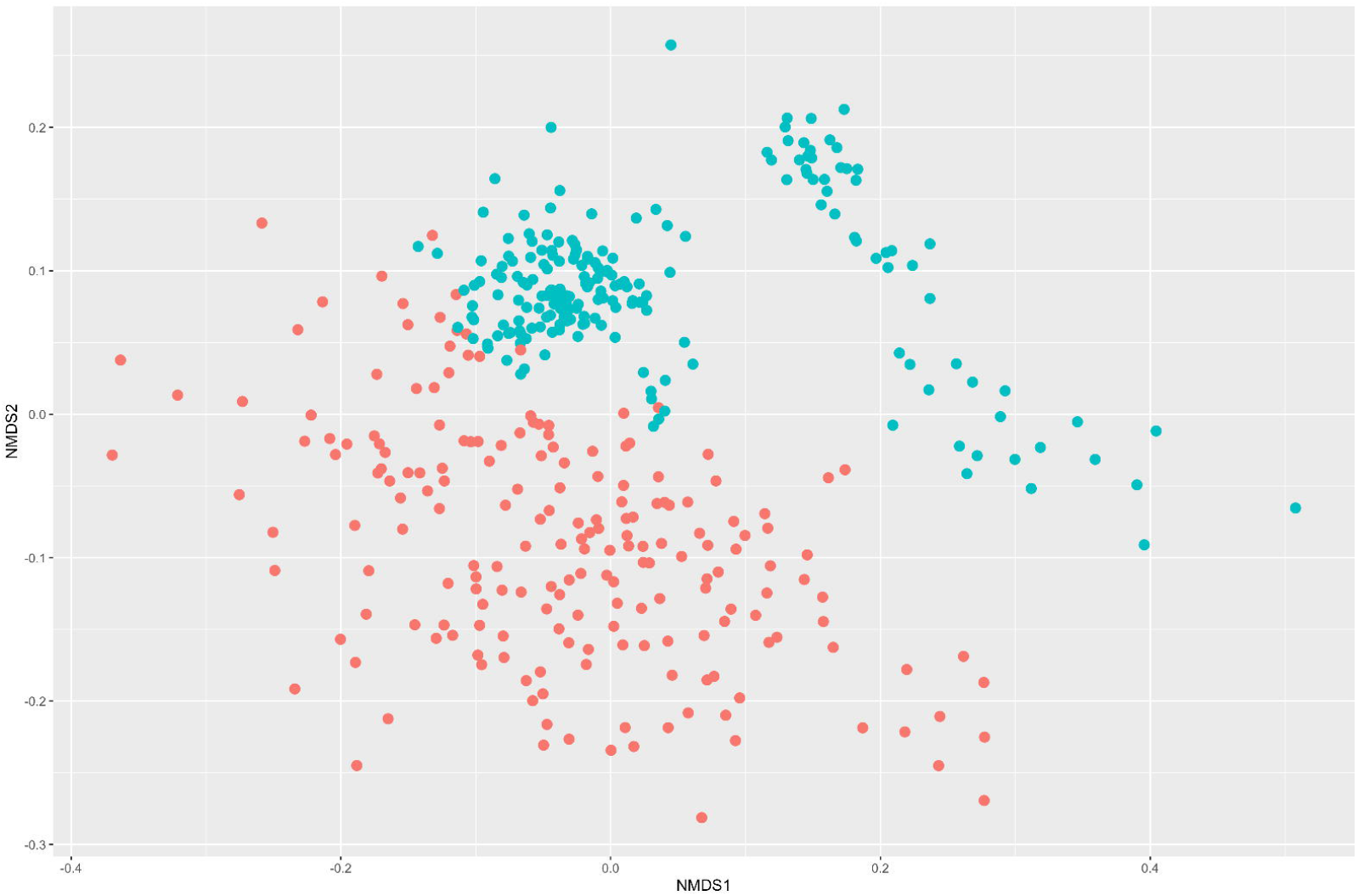
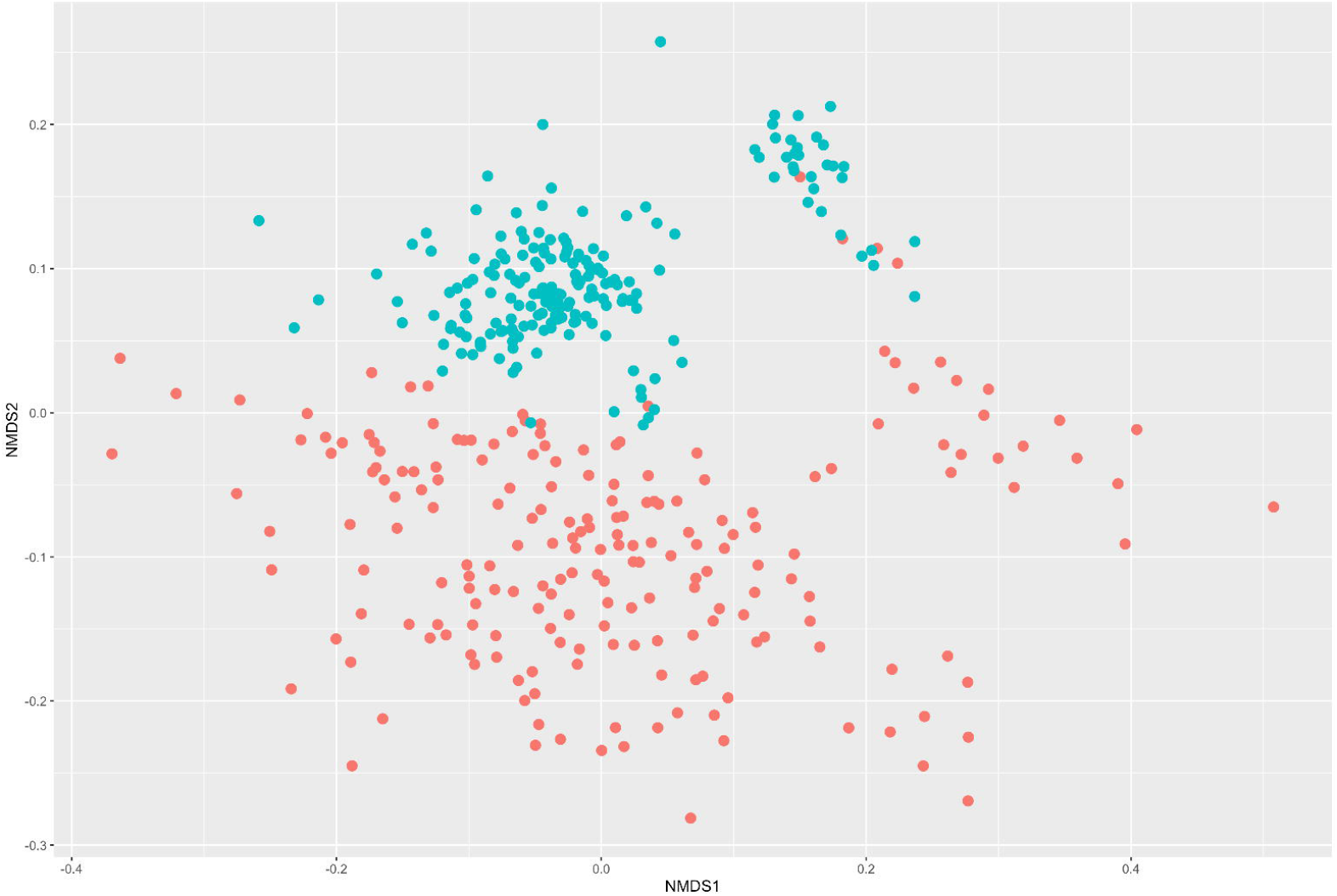
non-metric multidimensional scaling (NMDS) plot of dataset A with class labels predicted using the two approaches. **a.** Infinite Dirichlet multinomial mixture (DMM) with the stochastic variational variable selection (SVVS) algorithm; **b.** The finite DMM with EM algorithm *Note: Red-colored circles denote the control and blue-colored circles denote drought*

**Figure 2.**
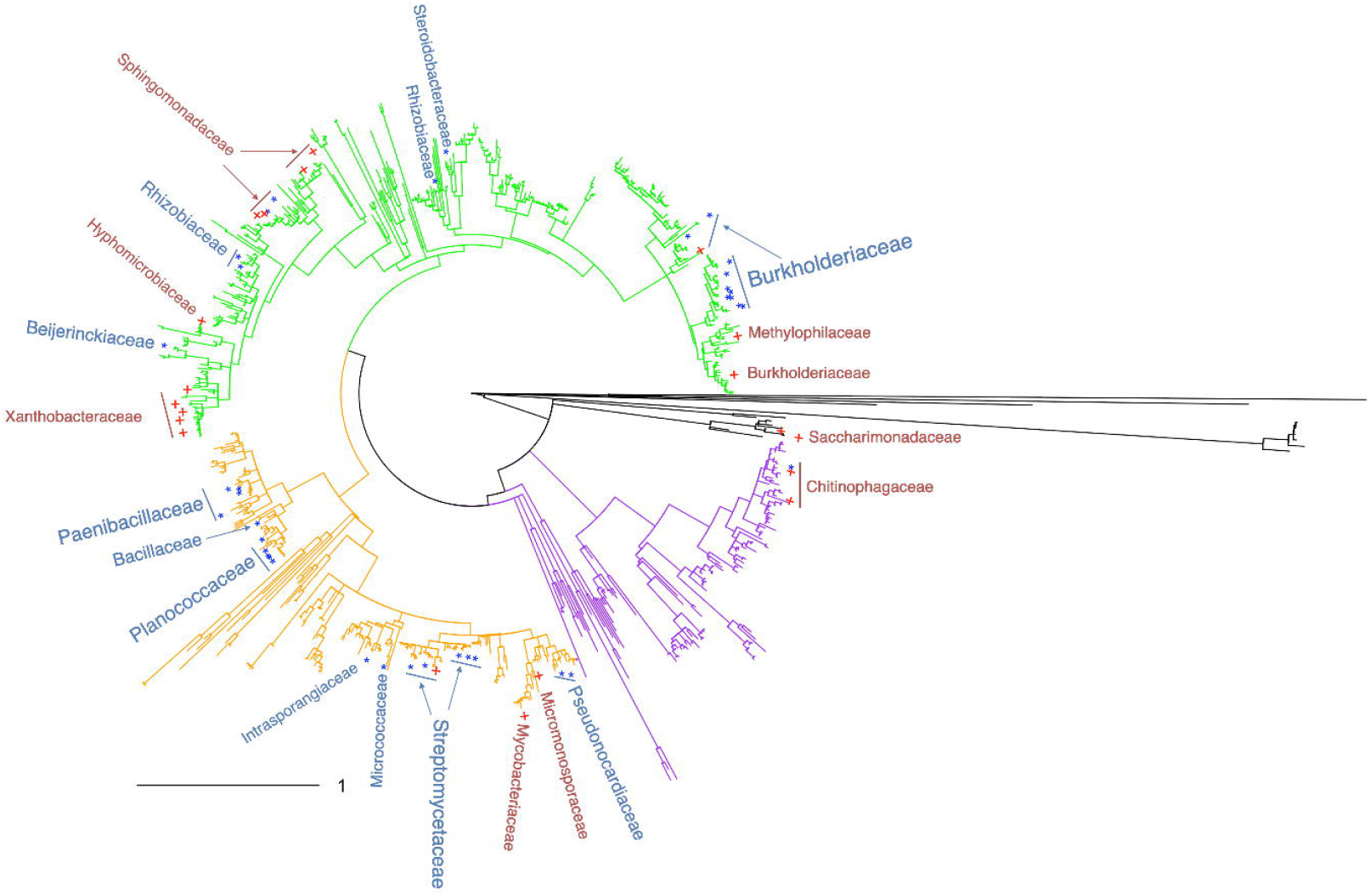
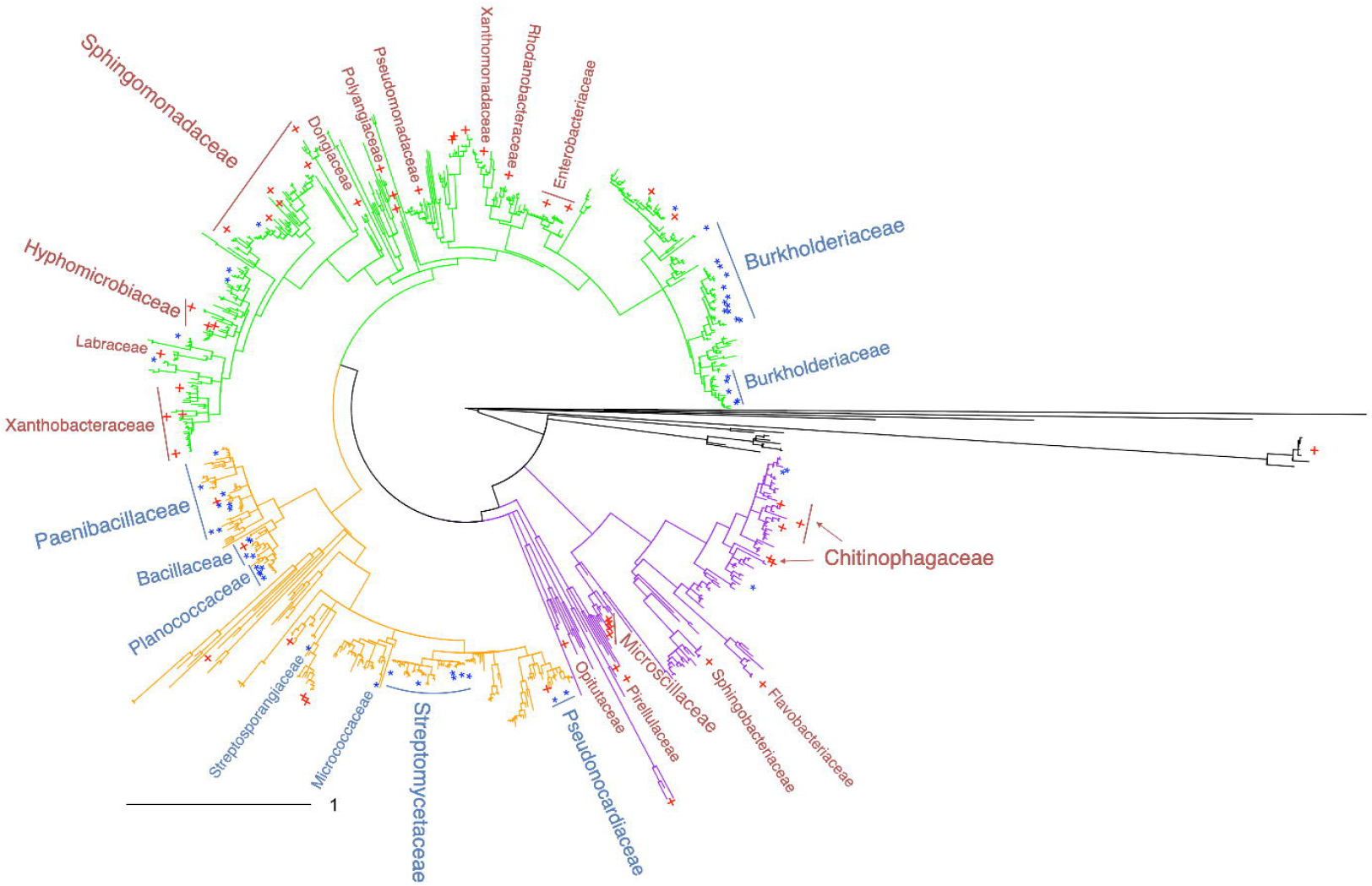
Microbial species selected using the stochastic variational variable selection (SVVS) approach and mapped on the phylogenetic tree for dataset A. **a.** Mapping of 50 selected microbiome species; **b.** Mapping of 100 selected microbiome species *Note: Red-colored plus symbols denote the control and blue-colored stars denote drought*

**Table 3.**
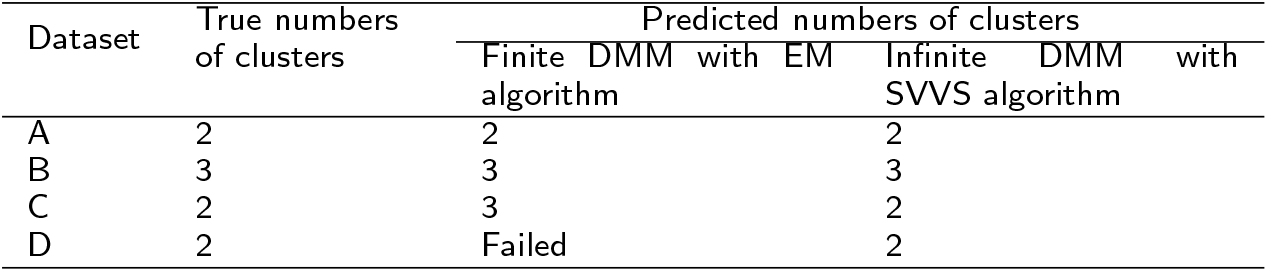
Numbers of clusters predicted by the two approaches for the real data sets

| Dataset | True numbers of clusters | Predicted numbers of clusters |  |
| --- | --- | --- | --- |
|  |  | Finite DMM with EM algorithm | Infinite DMM with SVVS algorithm |
| A | 2 | 2 | 2 |
| B | 3 | 3 | 3 |
| C | 2 | 3 | 2 |
| D | 2 | Failed | 2 |

**Table 4.** ARI scores of the two approaches for the real data sets

| Datasets | Finite DMM with EM algorithm | Infinite DMM with SVVS algorithm |
| --- | --- | --- |
| A | 0.76 | 0.98 |
| B | 0.44 | 0.66 |
| C | 0.21 | 0.48 |
| D | Failed | 0.5 |

### The selected microbiome species mapped on the phylogenetic trees

The other considerable contribution of SVVS is its ability to select a minimum core set of microbial species that shows significant differences among the clusters obtained in the analysis. Specifically, Figures 2a and b show that the top 50 and 100 selected microbiome species in dataset A were mapped on the 16S phylogenetic tree. Most microbiome families that were significantly associated with plant growth promotion under drought conditions were observed in the orange region of the tree. Our results are consistent with those of previous studies. For example, many species of bacterial families, including *Micrococcaceae, Paenibacillaceae, Bacillaceae,* and *Planococcaceae,* showed a strong dominance in ecosystems after the impact of wildfires on living organisms [47].

Moreover, Figure 3 shows that the top 100 selected microbiome species in dataset B were mapped on the 16S phylogenetic tree. The green region of the tree includes most microbiome species that show significant associations with non-diarrheal controls. Several dominant species that were significantly associated with CDI were observed in the orange region of the tree. A mixture of the two groups was observed in the purple region. Specifically, numerous microbiome species belonging to the *Bacteroidaceae, Porphyromonadaceae,* and *Rikenellaceae* families were observed in the green region of the tree. Several studies have shown that several bacterial species within these families are largely absent in CDI cases and are closely associated with non-diarrheal controls [25, 48]. One of the main risk factors is antibiotic treatments that alter the host nutritional landscape to produce the essential branched-chain amino acids and proline for *C. difficile* growth and to suppress the return of members of the *Rikenellaceae, Bacteroidaceae* families [49, 50].

**Figure 3.**
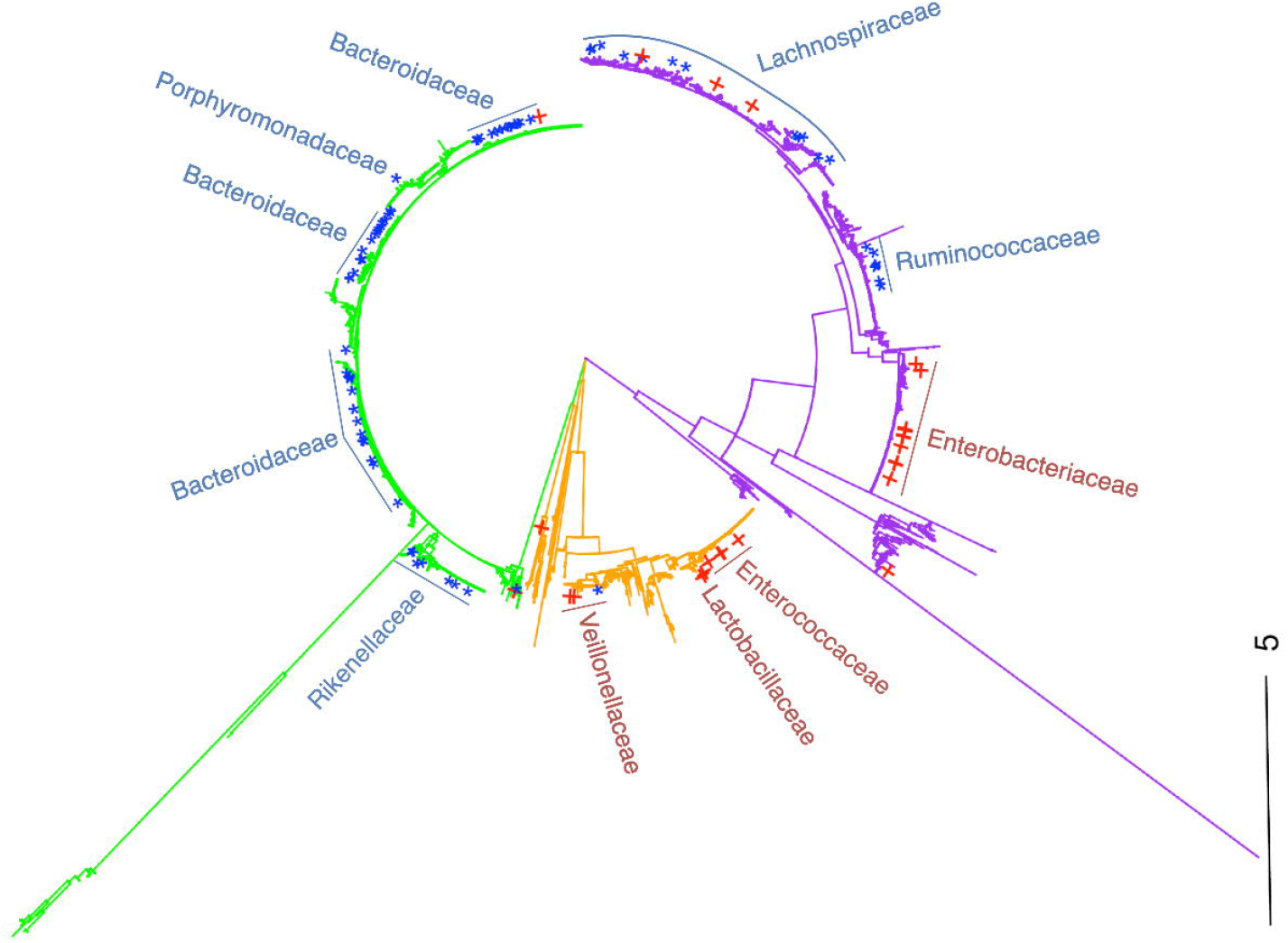
Microbial species selected using the stochastic variational variable selection (SVVS) approach and mapped on the phylogenetic tree for dataset B (*Clostridium difficile* infection (CDI) disease) *Note: Red-colored plus symbols denote for CDI cases and blue-colored stars denote non-diarrheal control*

### SVVS improves enterotype clustering

Figures 4a and b show that the SVVS algorithm of the infinite DMM model and the EM algorithm of the finite DMM model revealed two enterotypes of dataset E. The NMDS plots with the unweighted UniFrac distances showed that two enterotypes were clearly separated by the two approaches. Figure 4c shows the OTU-level Shannon diversity index was significantly different between the two enterotypes. Moreover, the top 100 selected microbiome species in dataset E, which significantly contributed to the enterotype clustering process, were mapped on the 16S phylogenetic tree. The Spearman correlations were applied to identify the 100 selected microbiome species associated with each enterotype. Figure 4d shows that the two enterotypes were clearly separated on the phylogenetic tree. Enterotype 2 had the highest levels of the genus *Bacteroides*. In the previous studies [51, 52, 53], the populations such as the European population, which consumes more animal protein and fats, show the dominance of the *Bacteroides* enterotype. Alternatively, Enterotype 1 showed a lower relative abundance of *Bacteroides* than in Enterotype 2 but had higher levels of the genera *Alistipes* and *Parabacteroides* (phylum *Bacteroidetes*), which characterize the *Bacteroides* enterotype. Moreover, the presence of the genera *Roseburia, Ruminococcus, Faecalibacterium, Subdoligranulum* and *Lachnospiraceae* (phylum *Firmicutes*) was observed in Enterotype 1.

**Figure 4.**
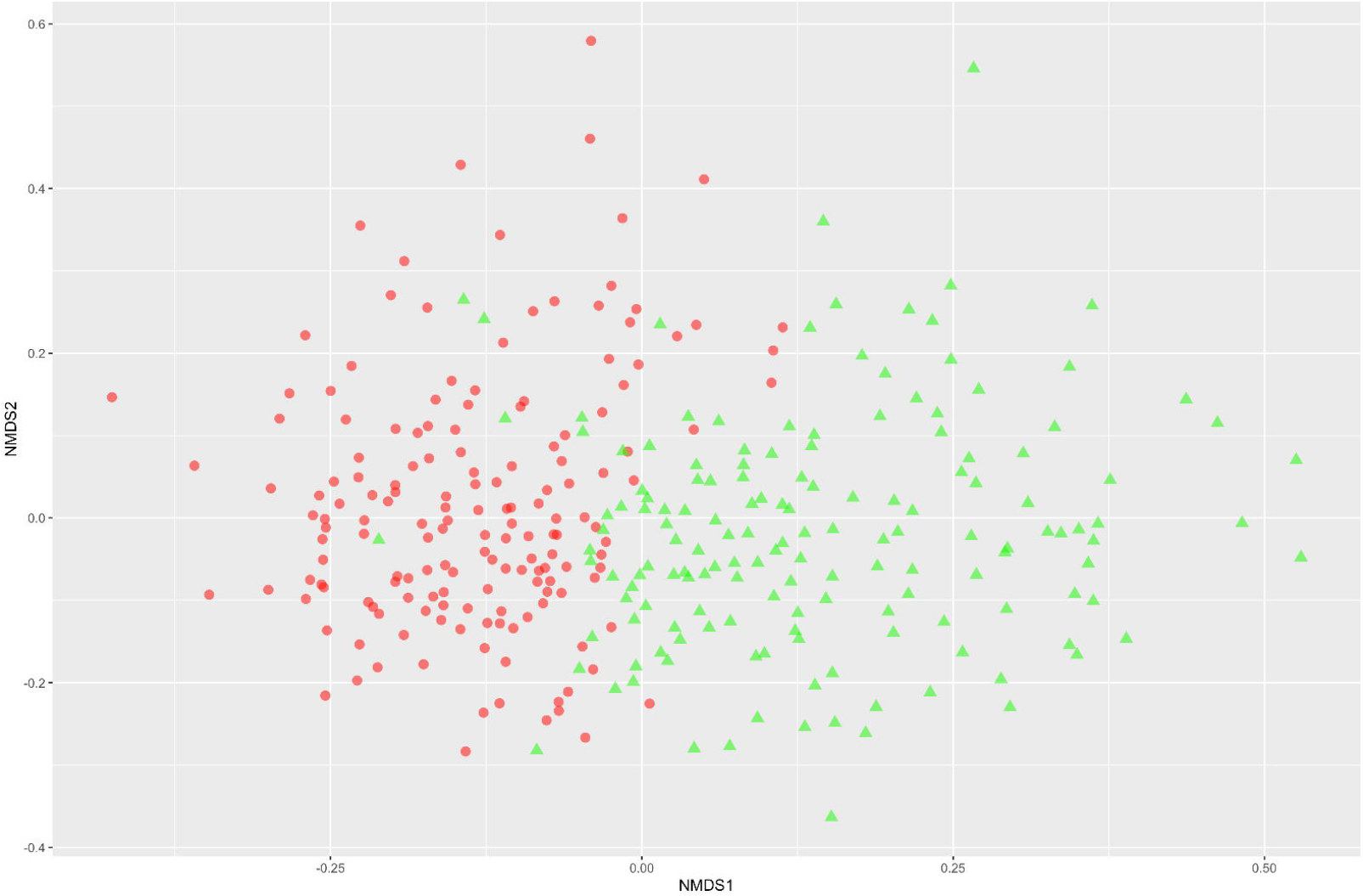
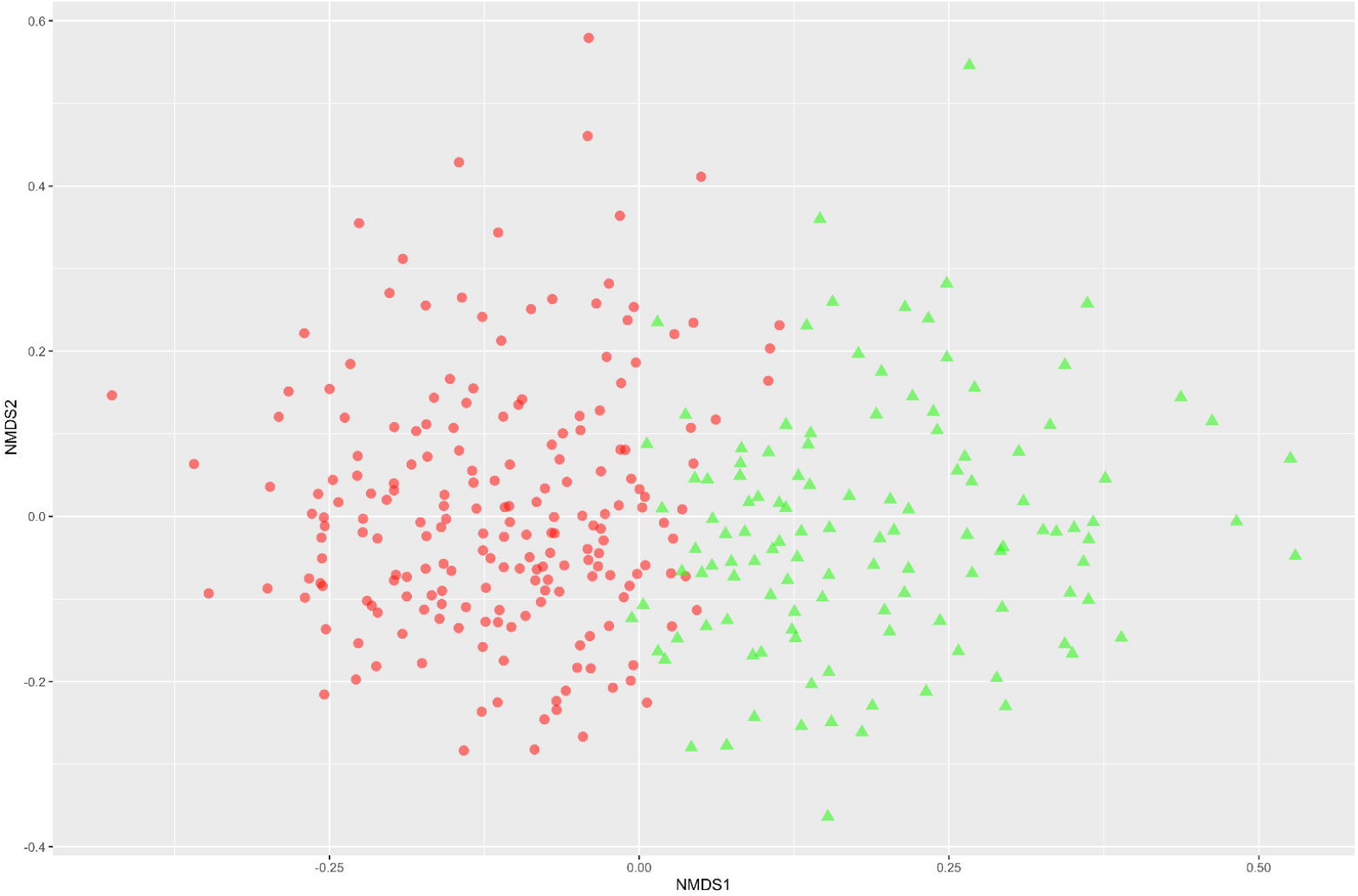
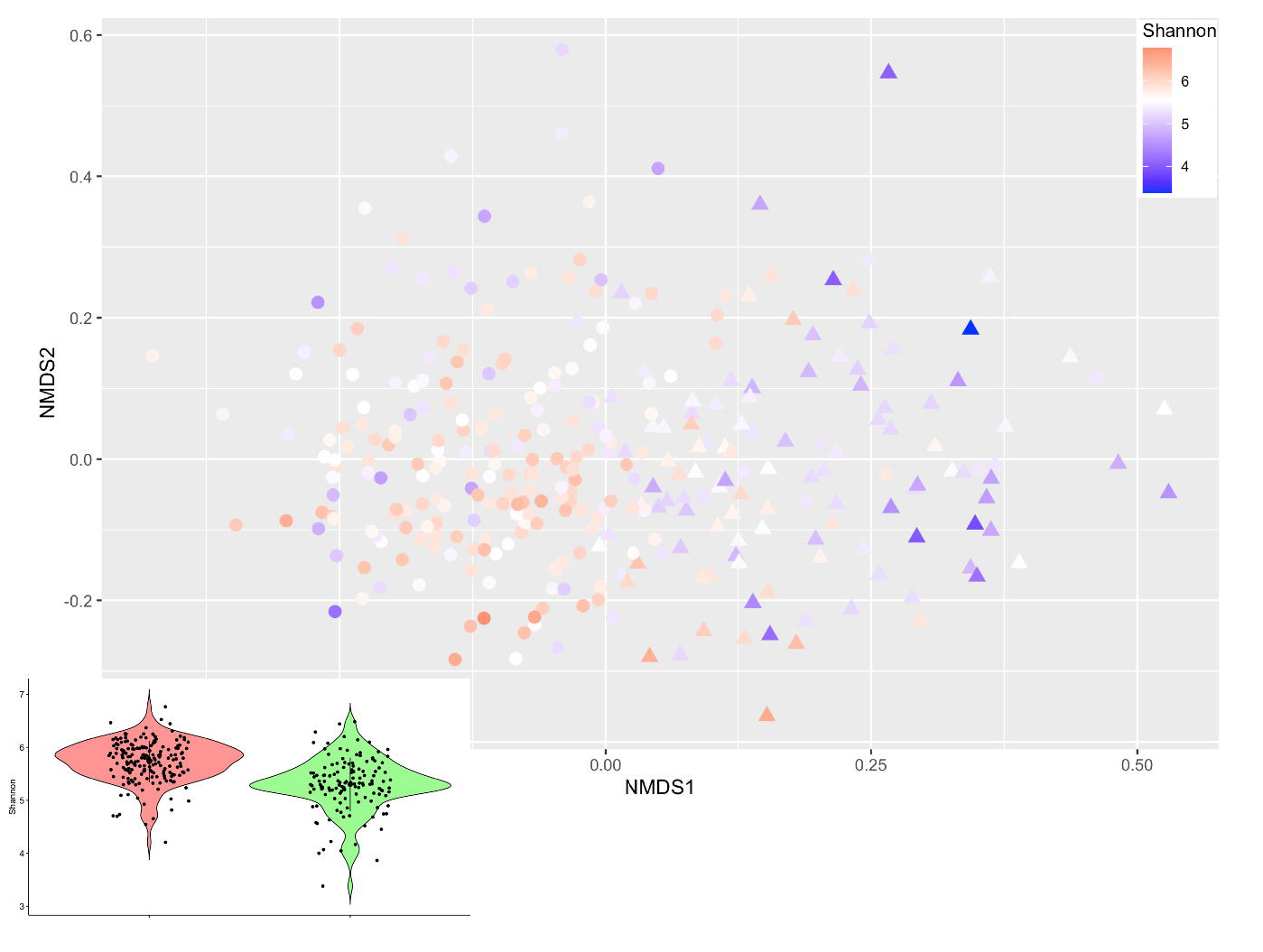
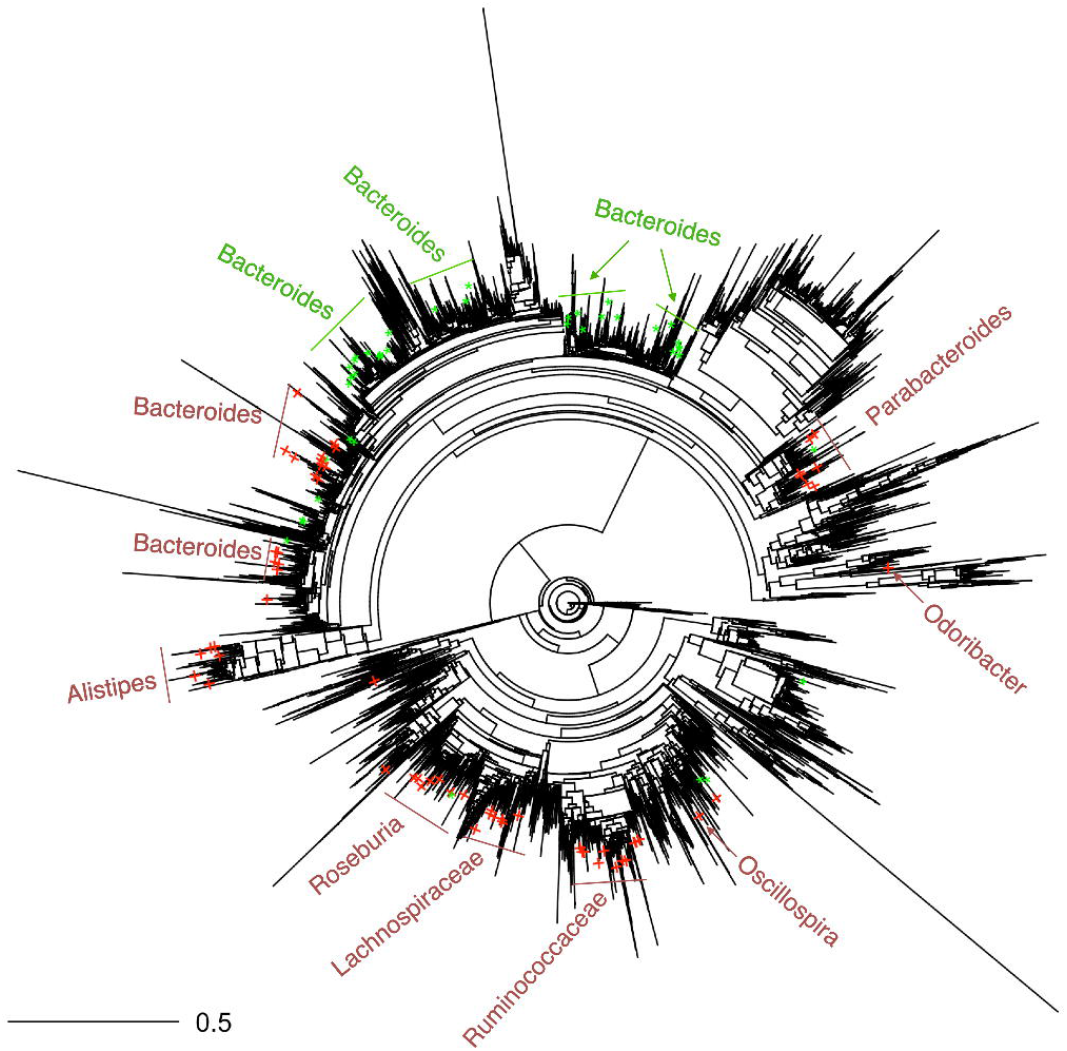
Enterotypes of the healthy human microbiome from the stool samples in dataset E. **a.** Labeled enterotypes of the finite Dirichlet multinomial mixture (DMM) with the EM algorithm; **b.** Labeled enterotypes of the infinite DMM with the stochastic variational variable selection (SVVS) algorithm; **c.** Alpha diversity of two enterotypes labeled by SVVS (Circle symbols denote Enterotype 1 and triangle symbols denote Enterotype 2); **d.** Top 100 important microbiome species that selected using the SVVS approach *Note: Red color denotes Enterotype 1 and green color denotes Enterotype 2*

## Discussion

Rapid identification of the minimum core set of OTUs in high-dimensional data of microbial studies is essential to further our understanding of microbial community structures in clustering analysis. The intensive concentration of a small number of relevant OTUs that significantly contribute to the task of clustering will not only increase the performance of these analyses but also open new opportunities for studies that explore the important associations of microbial communities with human diseases, precision medicine, and environmental conditions.

As the substantial increases in the dimensionality of the microbial datasets cause computational burden and poor performance with previous methods, the proposed approach could satisfy the high demands of the microbiome analysis. Our SVVS approach is useful in several aspects. First, SVVS integrated an indicator variable into the framework of the infinite DMM model to identify significant microbiome species (or OTUs) and used SVI to overcome computational limitations. Thus, the SVVS approach quickly identified the core set of microbial species (or OTUs), considerably improving the performance of the infinite DMM model. In particular, the SVVS method can complete its main tasks in massive microbiome datasets [26] that the previous methods could not perform. Second, a stick-breaking representation was proposed to extend the finite DMM model to an infinite case. This solution treated the total number of clusters as a free parameter of the model, which could help avoid the disadvantages of determining the number of clusters before running the algorithms. Therefore, SVVS could identify the main enterotypes of the healthy human microbiome and detect the important microbiome species that contribute to the variation of the different community compositions.

In recent years, several studies have highlighted the substantial role of large-scale analysis in discovering microbiome connections with host metabolism, host genetics in human health, medication [54, 55], and agroecosystems [38]. An increasing number of multi-omics datasets have been published, such as the integration of metagenomics, metatranscriptomics, metaproteomics [56], whole-genome sequencing, and whole-transcriptome sequencing of the TCGA cancer microbiome [57]. In the future, we plan to extend the SVVS approach to a comprehensive analysis of multi-omics datasets. The main framework of the SVVS method can be developed for the other Bayesian mixture models such as beta-mixture models for microarray gene expression datasets [58], and multinomial mixture model for ChIP-exo sequencing data [59]. Therefore, this method provides to new opportunities for discovering the significant associations of microbes with specific nutrients and medication or the important interactions between plants, microbes, and soils. Moreover, the dimensionality of databases significantly increased when other datasets were combined with microbiome data. We considered the integration of parallel computation strategies that can partition the data both horizontally (over sample) and vertically (over features) to speed up the computational processes [60].

## Conclusion

In conclude, the proposed stochastic variational variable selection approach can significantly improve the performance of the Dirichlet multinomial mixture model for analyzing high-dimensional microbial databases. The selected minimum core set of microbial species makes it easier to interpret the cause of sample differentiation. This study will contribute to and stimulate ongoing efforts to improve the performance of metagenomic statistical models that rapidly identify the potential species of the environmental and human microbiomes in a range of industrial sectors.

## Acknowledgements

We are grateful to the technical staff of the Arid Land Research Center, Tottori University, and Izumi Higashida for managing of the field experiments on soybean. We would like to thank all the members of the JST-CREST Program including Mikio Nakazono, Hirokazu Takahashi, Toru Fujiwara, Yoshihiro Ohmori, Hideki Takanashi, Akito Kaga, Mai Tsuda and Yuji Sawada for conducting the field experiments.

## Availability of data and materials

https://github.com/tungtokyo1108/SVVS

## Abbreviations

ARI: Adjusted Rand index
CDI: Clostridium difficile infection
DMM: Dirichlet multinomial mixture
EM: Expectation maximization
ELBO: Evidence lower bound
IBD: Inflammatory bowel disease
KL: Kullback-Leibler
MCMC: Markov chain Monte Carlo
OB: Obesity
QIIME: Quantitative Insights Into Microbial Ecology
OTUs: Operational taxonomic units
RI: Rand index
SVI: Stochastic variational inference
SVVS: Stochastic variational variable selection

## Ethics approval and consent to participate

Not applicable

## Consent for publication

Not applicable.

## Authors’ contributions

TD and HI designed the study. YT, YY, HT and HI designed and conducted the field experiment in Tottori. KK, EU, SK, TS and YI performed the microbiome analysis from tissue sampling, library preparation, sequencing, and primary informatics for taxonomic assignment and diversity statistics. TD developed the method and analyze the data. TD and HI interpreted the data and wrote the manuscript. The authors read and approved the final manuscript.

## Author details

## Tables

### Additional Files

Additional file 1: Supplemental Materials and Methods.

## References

1. Visconti, A., Le Roy, C.I., Rosa, F., Rossi, N., Martin, T.C., Mohney, R.P., Li, W., de Rinaldis, E., Bell, J.T., Venter, J.C., et al.: Interplay between the human gut microbiome and host metabolism. Nature communications 10(1), 1–10 (2019)

2. Al Nabhani, Z., Eberl, G.: Imprinting of the immune system by the microbiota early in life. Mucosal immunology 13(2), 183–189 (2020)

3. Emerson, J.B., Roux, S., Brum, J.R., Bolduc, B., Woodcroft, B.J., Jang, H.B., Singleton, C.M., Solden, L.M., Naas, A.E., Boyd, J.A., et al.: Host-linked soil viral ecology along a permafrost thaw gradient. Nature microbiology 3(8), 870–880 (2018)

4. Caporaso, J.G., Kuczynski, J., Stombaugh, J., Bittinger, K., Bushman, F.D., Costello, E.K., Fierer, N., Peña, A.G., Goodrich, J.K., Gordon, J.I., et al.: Qiime allows analysis of high-throughput community sequencing data. Nature methods 7(5), 335–336 (2010)

5. Holmes, I., Harris, K., Quince, C.: Dirichlet multinomial mixtures: generative models for microbial metagenomics. PloS one 7(2), 30126 (2012)

6. Depner, M., Taft, D.H., Kirjavainen, P.V., Kalanetra, K.M., Karvonen, A.M., Peschel, S., Schmausser-Hechfellner, E., Roduit, C., Frei, R., Lauener, R., et al.: Maturation of the gut microbiome during the first year of life contributes to the protective farm effect on childhood asthma. Nature Medicine 26(11), 1766–1775 (2020)

7. Vieira-Silva, S., Falony, G., Belda, E., Nielsen, T., Aron-Wisnewsky, J., Chakaroun, R., Forslund, S.K., Assmann, K., Valles-Colomer, M., Nguyen, T.T.D., et al.: Statin therapy is associated with lower prevalence of gut microbiota dysbiosis. Nature 581(7808), 310–315 (2020)

8. Hughes, D.A., Bacigalupe, R., Wang, J., Rühlemann, M.C., Tito, R.Y., Falony, G., Joossens, M., Vieira-Silva, S., Henckaerts, L., Rymenans, L., et al.: Genome-wide associations of human gut microbiome variation and implications for causal inference analyses. Nature microbiology 5(9), 1079–1087 (2020)

9. Zaneveld, J.R., McMinds, R., Thurber, R.V.: Stress and stability: applying the anna karenina principle to animal microbiomes. Nature microbiology 2(9), 1–8 (2017)

10. Papaspiliopoulos, O., Roberts, G.O.: Retrospective markov chain monte carlo methods for dirichlet process hierarchical models. Biometrika 95(1), 169–186 (2008)

11. Bouguila, N., Ziou, D.: A countably infinite mixture model for clustering and feature selection. Knowledge and information systems 33(2), 351–370 (2012)

12. Ferguson, T.S.: A bayesian analysis of some nonparametric problems. The annals of statistics, 209–230 (1973)

13. Green, P.J., Richardson, S.: Modelling heterogeneity with and without the dirichlet process. Scandinavian journal of statistics 28(2), 355–375 (2001)

14. Ishwaran, H., James, L.F.: Gibbs sampling methods for stick-breaking priors. Journal of the American Statistical Association 96(453), 161–173 (2001)

15. Jordan, M.I., Ghahramani, Z., Jaakkola, T.S., Saul, L.K.: An introduction to variational methods for graphical models. Machine learning 37(2), 183–233 (1999)

16. Blei, D.M., Jordan, M.I., et al.: Variational inference for dirichlet process mixtures. Bayesian analysis 1(1), 121–143 (2006)

17. Hoffman, M.D., Blei, D.M., Wang, C., Paisley, J.: Stochastic variational inference. Journal of Machine Learning Research 14(5) (2013)

18. Raj, A., Stephens, M., Pritchard, J.K.: faststructure: variational inference of population structure in large snp data sets. Genetics 197(2), 573–589 (2014)

19. Gopalan, P., Hao, W., Blei, D.M., Storey, J.D.: Scaling probabilistic models of genetic variation to millions of humans. Nature genetics 48(12), 1587 (2016)

20. Dang, T., Kishino, H.: Stochastic variational inference for bayesian phylogenetics: a case of cat model. Molecular biology and evolution 36(4), 825–833 (2019)

21. Fourment, M., Darling, A.E.: Evaluating probabilistic programming and fast variational bayesian inference in phylogenetics. PeerJ 7, 8272 (2019)

22. Fourment, M., Magee, A.F., Whidden, C., Bilge, A., Matsen IV, F.A., Minin, V.N.: 19 dubious ways to compute the marginal likelihood of a phylogenetic tree topology. Systematic biology 69(2), 209–220 (2020)

23. Ma, Z., Leijon, A.: Bayesian estimation of beta mixture models with variational inference. IEEE Transactions on Pattern Analysis and Machine Intelligence 33(11), 2160–2173 (2011)

24. Ma, Z., Rana, P.K., Taghia, J., Flierl, M., Leijon, A.: Bayesian estimation of dirichlet mixture model with variational inference. Pattern Recognition 47(9), 3143–3157 (2014)

25. Schubert, A.M., Rogers, M.A., Ring, C., Mogle, J., Petrosino, J.P., Young, V.B., Aronoff, D.M., Schloss, P.D.: Microbiome data distinguish patients with clostridium difficile infection and non-c. difficile-associated diarrhea from healthy controls. MBio 5(3), 01021–14 (2014)

26. Goodrich, J.K., Waters, J.L., Poole, A.C., Sutter, J.L., Koren, O., Blekhman, R., Beaumont, M., Van Treuren, W., Knight, R., Bell, J.T., et al.: Human genetics shape the gut microbiome. Cell 159(4), 789–799 (2014)

27. Gevers, D., Kugathasan, S., Denson, L.A., Vázquez-Baeza, Y., Van Treuren, W., Ren, B., Schwager, E., Knights, D., Song, S.J., Yassour, M., et al.: The treatment-naive microbiome in new-onset crohn’s disease. Cell host & microbe 15(3), 382–392 (2014)

28. Costea, P.I., Hildebrand, F., Arumugam, M., Bäckhed, F., Blaser, M.J., Bushman, F.D., De Vos, W.M., Ehrlich, S.D., Fraser, C.M., Hattori, M., et al.: Enterotypes in the landscape of gut microbial community composition. Nature microbiology 3(1), 8–16 (2018)

29. Schiffer, L., Azhar, R., Shepherd, L., Ramos, M., Geistlinger, L., Huttenhower, C., Dowd, J.B., Segata, N., Waldron, L.: Hmp16sdata: efficient access to the human microbiome project through bioconductor. American journal of epidemiology 188(6), 1023–1026 (2019)

30. Boutemedjet, S., Bouguila, N., Ziou, D.: A hybrid feature extraction selection approach for high-dimensional non-gaussian data clustering. IEEE Transactions on Pattern Analysis and Machine Intelligence 31(8), 1429–1443 (2008)

31. Dickey, J.M.: Multiple hypergeometric functions: Probabilistic interpretations and statistical uses. Journal of the American Statistical Association 78(383), 628–637 (1983)

32. Wang, C., Blei, D.M.: Variational inference in nonconjugate models. Journal of Machine Learning Research 14(4) (2013)

33. Amari, S.-I.: Differential geometry of curved exponential families-curvatures and information loss. The Annals of Statistics, 357–385 (1982)

34. Robbins, H., Monro, S.: A stochastic approximation method. The annals of mathematical statistics, 400–407 (1951)

35. Honkela, A., Raiko, T., Kuusela, M., Tornio, M., Karhunen, J.: Approximate riemannian conjugate gradient learning for fixed-form variational bayes. The Journal of Machine Learning Research 11, 3235–3268 (2010)

36. Rand, W.M.: Objective criteria for the evaluation of clustering methods. Journal of the American Statistical association 66(336), 846–850 (1971)

37. Kumaishi, K., Usui, E., Suzuki, K., Kobori, S., Sato, T., Toda, Y., Takanashi, H., Shinozaki, S., Noda, M., Takakura, A., et al.: Simple amplicon sequencing library preparation for plant root microbial community profiling. bioRxiv (2021)

38. Ichihashi, Y., Date, Y., Shino, A., Shimizu, T., Shibata, A., Kumaishi, K., Funahashi, F., Wakayama, K., Yamazaki, K., Umezawa, A., et al.: Multi-omics analysis on an agroecosystem reveals the significant role of organic nitrogen to increase agricultural crop yield. Proceedings of the National Academy of Sciences 117(25), 14552–14560 (2020)

39. Caporaso, J.G., Lauber, C.L., Walters, W.A., Berg-Lyons, D., Lozupone, C.A., Turnbaugh, P.J., Fierer, N., Knight, R.: Global patterns of 16s rrna diversity at a depth of millions of sequences per sample. Proceedings of the national academy of sciences 108(Supplement 1), 4516–4522 (2011)

40. Lundberg, D.S., Yourstone, S., Mieczkowski, P., Jones, C.D., Dangl, J.L.: Practical innovations for high-throughput amplicon sequencing. Nature methods 10(10), 999–1002 (2013)

41. Callahan, B.J., McMurdie, P.J., Rosen, M.J., Han, A.W., Johnson, A.J.A., Holmes, S.P.: Dada2: high-resolution sample inference from illumina amplicon data. Nature methods 13(7), 581–583 (2016)

42. Quast, C., Pruesse, E., Yilmaz, P., Gerken, J., Schweer, T., Yarza, P., Peplies, J., Glöckner, F.O.: The silva ribosomal rna gene database project: improved data processing and web-based tools. Nucleic acids research 41(D1), 590–596 (2012)

43. Oksanen, J., Blanchet, F., Friendly, M., Kindt, R., Legendre, P., McGlinn, D., et al.: vegan: community ecology package. R package version 2.5-7. 2020

44. Duvallet, C., Gibbons, S.M., Gurry, T., Irizarry, R.A., Alm, E.J.: Meta-analysis of gut microbiome studies identifies disease-specific and shared responses. Nature communications 8(1), 1–10 (2017)

45. Morgan, M.: Dirichletmultinomial: Dirichlet-multinomial mixture model machine learning for microbiome data. R package version 1.34.0 1(0) (2021)

46. McMurdie, P.J., Holmes, S.: phyloseq: an r package for reproducible interactive analysis and graphics of microbiome census data. PloS one 8(4), 61217 (2013)

47. de Miera, L.E.S., Pinto, R., Gutierrez-Gonzalez, J.J., Calvo, L., Ansola, G.: Wildfire effects on diversity and composition in soil bacterial communities. Science of the Total Environment 726, 138636 (2020)

48. Rousseau, C., Poilane, I., De Pontual, L., Maherault, A.-C., Le Monnier, A., Collignon, A.: Clostridium difficile carriage in healthy infants in the community: a potential reservoir for pathogenic strains. Clinical Infectious Diseases 55(9), 1209–1215 (2012)

49. Hofmann, J.D., Otto, A., Berges, M., Biedendieck, R., Michel, A.-M., Becher, D., Jahn, D., Neumann-Schaal, M.: Metabolic reprogramming of clostridioides difficile during the stationary phase with the induction of toxin production. Frontiers in microbiology 9, 1970 (2018)

50. Fletcher, J.R., Pike, C.M., Parsons, R.J., Rivera, A.J., Foley, M.H., McLaren, M.R., Montgomery, S.A., Theriot, C.M.: Clostridioides difficile exploits toxin-mediated inflammation to alter the host nutritional landscape and exclude competitors from the gut microbiota. Nature Communications 12(1), 1–14 (2021)

51. De Filippo, C., Cavalieri, D., Di Paola, M., Ramazzotti, M., Poullet, J.B., Massart, S., Collini, S., Pieraccini, G., Lionetti, P.: Impact of diet in shaping gut microbiota revealed by a comparative study in children from europe and rural africa. Proceedings of the National Academy of Sciences 107(33), 14691–14696 (2010)

52. Wu, G.D., Chen, J., Hoffmann, C., Bittinger, K., Chen, Y.-Y., Keilbaugh, S.A., Bewtra, M., Knights, D., Walters, W.A., Knight, R., et al.: Linking long-term dietary patterns with gut microbial enterotypes. Science 334(6052), 105–108 (2011)

53. Mobeen, F., Sharma, V., Tulika, P.: Enterotype variations of the healthy human gut microbiome in different geographical regions. Bioinformation 14(9), 560 (2018)

54. Kurilshikov, A., Medina-Gomez, C., Bacigalupe, R., Radjabzadeh, D., Wang, J., Demirkan, A., Le Roy, C.I., Garay, J.A.R., Finnicum, C.T., Liu, X., et al.: Large-scale association analyses identify host factors influencing human gut microbiome composition. Nature Genetics 53(2), 156–165 (2021)

55. Asnicar, F., Berry, S.E., Valdes, A.M., Nguyen, L.H., Piccinno, G., Drew, D.A., Leeming, E., Gibson, R., Le Roy, C., Al Khatib, H., et al.: Microbiome connections with host metabolism and habitual diet from 1,098 deeply phenotyped individuals. Nature Medicine 27(2), 321–332 (2021)

56. Hultman, J., Waldrop, M.P., Mackelprang, R., David, M.M., McFarland, J., Blazewicz, S.J., Harden, J., Turetsky, M.R., McGuire, A.D., Shah, M.B., et al.: Multi-omics of permafrost, active layer and thermokarst bog soil microbiomes. Nature 521(7551), 208–212 (2015)

57. Poore, G.D., Kopylova, E., Zhu, Q., Carpenter, C., Fraraccio, S., Wandro, S., Kosciolek, T., Janssen, S., Metcalf, J., Song, S.J., et al.: Microbiome analyses of blood and tissues suggest cancer diagnostic approach. Nature 579(7800), 567–574 (2020)

58. Ji, Y., Wu, C., Liu, P., Wang, J., Coombes, K.R.: Applications of beta-mixture models in bioinformatics. Bioinformatics 21(9), 2118–2122 (2005)

59. Yamada, N., Lai, W.K., Farrell, N., Pugh, B.F., Mahony, S.: Characterizing protein–dna binding event subtypes in chip-exo data. Bioinformatics 35(6), 903–913 (2019)

60. Xing, E.P., Ho, Q., Xie, P., Wei, D.: Strategies and principles of distributed machine learning on big data. Engineering 2(2), 179–195 (2016)

