## Supplementary material for "Stochastic variational variable selection for high-dimensional microbiome data": Supp

#### 1 Mean-Field Variational Inference for DMM Model

We expand specifically the variational lower bound equation as follows:

$$\begin{aligned}\mathcal{L} &= \sum_{k=1}^{K_{max}} \mathrm{E}_q\left[\log\left(p\left(\gamma_k|1,\kappa\right)\right)\right] - \sum_{k=1}^{K_{\max}} \mathrm{E}_q\left[\log\left(q\left(\gamma_k|\vartheta_k,\vartheta_k^{'}\right)\right)\right] \\ &+ \sum_{i=1}^{N}\sum_{k=1}^{K_{max}} \mathrm{E}_q\left[\log p(Z_i^k|\gamma_1,\gamma_2,...,\gamma_{K_{max}})\right] - \sum_{i=1}^{N}\sum_{k=1}^{K_{\max}} \mathrm{E}_q\left[\log\left(q\left(Z_i^k|r_i^k\right)\right)\right] \\ &+ \sum_{i=1}^{N}\sum_{j=1}^{S} \mathrm{E}_q\left[\log p(\phi_{ij}|\epsilon_{j_1},\epsilon_{j_2})\right] - \sum_{i=1}^{N}\sum_{j=1}^{S} \mathrm{E}_q\left[\log\left(q\left(\phi_{ij}|f_{ij}\right)\right)\right] \\ &+ \sum_{j}^{S} \mathrm{E}_q\left[\log\left(p\left(\epsilon_j|\xi\right)\right)\right] - \sum_{j}^{S} \mathrm{E}_q\left[\log\left(q\left(\epsilon_j|\xi^*\right)\right)\right] \\ &+ \sum_{j=1}^{S}\sum_{k=1}^{K_{max}} \mathrm{E}_q\left[\log\left(p\left(\alpha_{kj}|\lambda_{kj}\right)\right)\right] - \sum_{j=1}^{S}\sum_{k=1}^{K_{max}} \mathrm{E}_q\left[\log\left(q\left(\alpha_{kj}|\lambda_{kj}^*\right)\right)\right] \\ &+ \sum_{j=1}^{S} \mathrm{E}_q\left[\log\left(p\left(\beta_j|\iota_j\right)\right)\right] - \sum_{j=1}^{S} \mathrm{E}_q\left[\log\left(q\left(\beta_j|\iota_j^*\right)\right)\right]\end{aligned} \tag{1}$$

To compute the variational expectations $E_q[.]$ in equation (1), we use the properties of exponential family distribution. If variational distributions for $q(\alpha_{kj}|\lambda_{kj})$ and $q(\beta_j|\iota_j)$ are Dirichlet distributions, then the exponential family representations are given by

$$\begin{aligned} q\left(\boldsymbol{\alpha}_k|\boldsymbol{\lambda}_k\right) &= \exp\left[\left(\sum_{j=1}^{S}\left(\lambda_j^k-1\right)\log\left(\alpha_j^k\right)\right) + \log\Gamma\left(\sum_{j=1}^{S}\lambda_j^k\right) - \sum_{j=1}^{S}\left(\log\Gamma\left(\lambda_j^k\right)\right)\right] \\ q\left(\boldsymbol{\beta}|\boldsymbol{\iota}\right) &= \exp\left[\left(\sum_{j=1}^{S}\left(\iota_j-1\right)\log\left(\beta_j\right)\right) + \log\Gamma\left(\sum_{j=1}^{S}\iota_j\right) - \sum_{j=1}^{S}\left(\log\Gamma\left(\iota_j\right)\right)\right]\end{aligned}$$

So the natural parameters and sufficient statistics of the Dirichlet distributions for $\alpha$ and $\beta$ are $\eta_{\alpha_{kj}} = \lambda_{kj}-1$, $T(\alpha_{kj}) = \log(\alpha_{kj})$ and $\eta_{\beta_j} = \iota_j - 1$, $T(\beta_j) = \log(\beta_j)$, resepctively. The approximation expectations are

$$\mathrm{E}\left[\alpha_{kj}\right] = \frac{\lambda_{kj}}{\sum_{j'=1}^{S}\lambda_{kj'}} \qquad \mathrm{E}\left[\beta_j\right] = \frac{\iota_j}{\sum_{j'=1}^{S}\iota_{j'}}$$

$$\mathrm{E}\left[\log\left(\alpha_{kj}\right)\right] = \psi\left(\lambda_{kj}\right) - \psi\left(\sum_{j'=1}^{S}\lambda_{kj'}\right) \qquad \mathrm{E}\left[\log\left(\beta_j\right)\right] = \psi\left(\iota_j\right) - \psi\left(\sum_{j'=1}^{S}\iota_{j'}\right)$$

where $\psi(.)$ is the digamma function.

With truncated stick-breaking representation of Dirichlet Mixture process, variational factors for the stick lengths $q\left(\gamma_k|\vartheta_k,\vartheta_k^{'}\right)$ are Beta distributions and the assignment variable $Z_i^k = \mathrm{I}\left[Z_i = k\right]$ for the $i^{th}$ sample allocation is governed by a multinomial distribution indexed by a variational parameter $r_{ik}$. The computation of the approximate expectation for the parameters of truncated stick-breaking representation have been considered carefully by Blei and Jordan (2006). The results of variational expectation for both variables are as follows:

$$
\begin{aligned}
&q\left(Z_i = k\right) = r_i^k \\
&q\left(Z_i > k\right) = \sum_{k'=k+1}^{K_{\max}} r_i^{k'} \\
&\mathrm{E}_q\left[\log\left(\gamma_k\right)\right] = \psi\left(\vartheta_k\right) - \psi\left(\vartheta_k + \vartheta_k^{'}\right) \\
&\mathrm{E}_q\left[\log\left(1-\gamma_k\right)\right] = \psi\left(\vartheta_k^{'}\right) - \psi\left(\vartheta_k + \vartheta_k^{'}\right)
\end{aligned}
$$

### 2 Stochastic Optimization of the Variational Parameters

#### 2.1 Compute the variational function parameters for the position of sample $i$ to be reallocated to cluster $k$

Following the principles of the variational inference (Bishop. 2006, Blei and Jordan. 2006, Hoffman 2013), variational parameters of $Z_i^k$ is the local parameters, we consider the derivation of the update equation for only one variable by fixing the others' distribution. The optimal coordinate updates are derived for local parameters. The log of the optimized factor to the posterior distribution of $Z_i^k$ is

$$
\log Q^*\left(Z\right) = E_{\phi,\gamma,\epsilon,\alpha,\beta}\left[\log p\left(Z,\phi,\gamma,\epsilon,\alpha,\beta\right)\right] + const = \sum_{i=1}^{N}\sum_{k=1}^{K} Z_i^k \log\left(r_i^k\right) + const
$$

$$
\begin{aligned}
\log r_i^k = &\textstyle\sum_{j=1}^S \mathrm{E}_Q\left[\phi_{ij}\right]\left(\mathrm{E}_Q\left[log\left(\frac{\Gamma(\sum_{j=1}^S \alpha_{kj})}{\Gamma(\sum_{j=1}^S X_{ij}+\sum_{j=1}^S \alpha_{kj})}\right)\right] + \sum_{j=1}^S \mathrm{E}_Q\left[log\left(\frac{\Gamma(X_{ij}+\alpha_{kj})}{\Gamma(\alpha_{kj})}\right)\right] + log(J_i!) + \sum_{j=1}^S log\left(\frac{1}{X_{ij}!}\right)\right) \\
&+\mathrm{E}_Q\left[log(\lambda_k)\right] + \textstyle\sum_{k'=1}^{k-1}\mathrm{E}_Q\left[log(1-\lambda_{k'})\right]
\end{aligned} \tag{2}
$$

$$
\mathrm{E}_Q\left[Z_i^k\right] = r_i^k = \exp\left\{\log\left(r_i^k\right)\right\}
$$

The expected logarithm of the functions $\mathrm{E}_Q\left[log\left(\frac{\Gamma(\sum_{j=1}^S \alpha_{kj})}{\Gamma(\sum_{j=1}^S X_{ij}+\sum_{j=1}^S \alpha_{kj})}\right)\right]$ and $\mathrm{E}_Q\left[log\left(\frac{\Gamma(X_{ij}+\alpha_{kj})}{\Gamma(\alpha_{kj})}\right)\right]$ in equation (2) have not the closed form. Thus, the calculation of these equations are analytically intractable. To use standard form of the variational inference (Bishop. 2006) or the stochastic variational inference (Hoffman et al. 2013), we need a closed-form expression. In order to overcome this problem, we consider a common approach:

- By applying a first-order Taylor expansion to preserve a bound, intractable expectations are avoided. By using these results, the variational parameters are updated as follow:

$$
\begin{aligned}
&log\left(\frac{\Gamma(X_{ij}+\alpha_{kj})}{\Gamma(\alpha_{kj})}\right) \geq log\left(\frac{\Gamma(X_{ij}+\overline{\alpha_{kj}})}{\Gamma(\overline{\alpha_{kj}})}\right) + \frac{\partial F(\alpha_{kj})}{\partial \alpha_{kj}}\frac{\partial \alpha_{kj}}{\partial log(\alpha_{kj})}|_{\alpha_{kj}=\overline{\alpha_{kj}}}\left(log(\alpha_{kj}) - log(\overline{\alpha_{kj}})\right) \\
&\mathrm{E}_Q\left[log\left(\frac{\Gamma(X_{ij}+\alpha_{kj})}{\Gamma(\alpha_{kj})}\right)\right] \geq log\left(\frac{\Gamma(X_{ij}+\overline{\alpha_{kj}})}{\Gamma(\overline{\alpha_{kj}})}\right) + \overline{\alpha_{kj}}\left[\Psi\left(\overline{\alpha_{kj}}+X_{ij}\right) - \Psi\left(\overline{\alpha_{kj}}\right)\right]\left(\mathrm{E}_Q\left[log(\alpha_{kj})\right] - log(\overline{\alpha_{kj}})\right) \\
&\geq log\left(\frac{\Gamma(X_{ij}+\overline{\alpha_{kj}})}{\Gamma(\overline{\alpha_{kj}})}\right) + \overline{\alpha_{kj}}\left[\Psi\left(\overline{\alpha_{kj}}+X_{ij}\right) - \Psi\left(\overline{\alpha_{kj}}\right)\right]\left(\Psi(\lambda_{kj}^*) - \Psi\left(\textstyle\sum_{j=1}^S \lambda_{kj}^*\right) - log(\overline{\alpha_{kj}})\right)
\end{aligned}
$$

$$
\begin{aligned}
&\mathrm{E}_Q\left[log\left(\frac{\Gamma(\sum_{j=1}^S \alpha_{kj})}{\Gamma(\sum_{j=1}^S X_{ij}+\sum_{j=1}^S \alpha_{kj})}\right)\right] \geq log\left(\frac{\Gamma(\sum_{j=1}^S \overline{\alpha_{kj}})}{\Gamma(\sum_{j=1}^S X_{ij}+\sum_{j=1}^S \overline{\alpha_{kj''}})}\right) \\
&+\textstyle\sum_{j=1}^S\left[\Psi\left(\sum_{j=1}^S \overline{\alpha_{kj}}\right) - \Psi\left(\sum_{j=1}^S X_{ij} + \sum_{j=1}^S \overline{\alpha_{kj}}\right)\right]\overline{\alpha_{kj}}\left(\Psi(\lambda_{kj}^*) - \Psi\left(\sum_{j=1}^S \lambda_{kj}^*\right) - log(\overline{\alpha_{kj}})\right)
\end{aligned}
$$

Based on this framework, the variational parameters of $Z_{ik}$ are updated as follow:

$$
\begin{aligned}
\log r_i^k = &\textstyle\sum_{j=1}^S f_{ij} log\left(\frac{\Gamma(\sum_{j=1}^S \overline{\alpha_{kj}})}{\Gamma(\sum_{j=1}^S X_{ij}+\sum_{j=1}^S \overline{\alpha_{kj''}})}\right) \\
&+\textstyle\sum_{j=1}^S f_{ij}\left[\Psi\left(\sum_{j=1}^S \overline{\alpha_{kj}}\right) - \Psi\left(\sum_{j=1}^S X_{ij} + \sum_{j=1}^S \overline{\alpha_{kj}}\right)\right]\overline{\alpha_{kj}}\left(\Psi(\lambda_{kj}^*) - \Psi\left(\sum_{j=1}^S \lambda_{kj}^*\right) - log(\overline{\alpha_{kj}})\right) \\
&+\textstyle\sum_{j=1}^S f_{ij}\left[log\left(\frac{\Gamma(X_{ij}+\overline{\alpha_{kj}})}{\Gamma(\overline{\alpha_{kj}})}\right) + \overline{\alpha_{kj}}\left[\Psi\left(\overline{\alpha_{kj}}+X_{ij}\right) - \Psi\left(\overline{\alpha_{kj}}\right)\right]\left(\Psi(\lambda_{kj}^*) - \Psi\left(\sum_{j=1}^S \lambda_{kj}^*\right) - log(\overline{\alpha_{kj}})\right)\right] \\
&+\Psi(\vartheta_k) - \Psi(\vartheta_k + \vartheta_k^{'}) + \textstyle\sum_{k'=1}^{k-1}\Psi(\vartheta_{k'}^{'}) - \Psi(\vartheta_{k'} + \vartheta_{k'}^{'})
\end{aligned} \tag{3}
$$

#### 2.2 Compute the variational function parameters of indicator variables for microbiome selection

Similarity, the optimized solution to the posterior distribution of variable selection $\phi_{ij}$

$$\log Q^*(\phi) = E_{Z,\gamma,\epsilon,\alpha,\beta}\left[\log p(Z,\phi,\gamma,\epsilon,\alpha,\beta)\right] + const = \sum_{i=1}^{N}\sum_{j=1}^{S}\phi_{ij}\log(f_{ij}) + const$$

$$\begin{aligned}\log f_{ij}^{\phi_{ij}} &= \textstyle\sum_{k=1}^{K_{max}} r_{ik} log\left(\frac{\Gamma(\sum_{j=1}^{S}\overline{\alpha_{kj}})}{\Gamma(\sum_{j=1}^{S}X_{ij}+\sum_{j=1}^{S}\overline{\alpha_{kj''}})}\right)\\ &+\textstyle\sum_{k=1}^{K_{max}} r_{ik}\left[\Psi\left(\sum_{j=1}^{S}\overline{\alpha_{kj}}\right)-\Psi\left(\sum_{j=1}^{S}X_{ij}+\sum_{j=1}^{S}\overline{\alpha_{kj}}\right)\right]\overline{\alpha_{kj}}\left(\Psi(\lambda_{kj}^*)-\Psi\left(\sum_{j=1}^{S}\lambda_{kj}^*\right)-log(\overline{\alpha_{kj}})\right)\\ &+\textstyle\sum_{k=1}^{K_{max}} r_{ik}\left[log\left(\frac{\Gamma(X_{ij}+\overline{\alpha_{kj}})}{\Gamma(\overline{\alpha_{kj}})}\right)+\overline{\alpha_{kj}}\left[\Psi\left(\overline{\alpha_{kj}}+X_{ij}\right)-\Psi\left(\overline{\alpha_{kj}}\right)\right]\left(\Psi(\lambda_{kj}^*)-\Psi\left(\sum_{j=1}^{S}\lambda_{kj}^*\right)-log(\overline{\alpha_{kj}})\right)\right]\\ &+[\Psi(\xi_1^*)-\Psi(\xi_1^*+\xi_2^*)]\end{aligned} \quad (4)$$

$$\begin{aligned}\log f_{ij}^{1-\phi_{ij}} &= log\left(\frac{\Gamma(\sum_{j=1}^{S}\overline{\beta_j})}{\Gamma(\sum_{j=1}^{S}X_{ij}+\sum_{j=1}^{S}\overline{\beta_{j''}})}\right)\\ &+\textstyle\sum_{j=1}^{S}\left[\Psi\left(\sum_{j=1}^{S}\overline{\beta_j}\right)-\Psi\left(\sum_{j=1}^{S}X_{ij}+\sum_{j=1}^{S}\overline{\beta_j}\right)\right]\overline{\beta_j}\left(\Psi(\iota_j^*)-\Psi\left(\sum_{j=1}^{S}\iota_j^*\right)-log(\overline{\beta_j})\right)\\ &+log\left(\frac{\Gamma(X_{ij}+\overline{\beta_j})}{\Gamma(\overline{\beta_j})}\right)+\overline{\beta_j}\left[\Psi\left(\overline{\beta_j}+X_{ij}\right)-\Psi\left(\overline{\beta_j}\right)\right]\left(\Psi(\iota_j^*)-\Psi\left(\sum_{j=1}^{S}\iota_j^*\right)-log(\overline{\alpha_{kj}})\right)\\ &+[\Psi(\xi_2^*)-\Psi(\xi_1^*+\xi_2^*)]\end{aligned} \quad (5)$$

#### 2.3 Updating variational parameters of $\alpha$

Based on the principal framework of the stochastic variational inference (Hoffman et al. 2013), we consider parameters of $\alpha$ as global parameters. These parameters are updated by a stochastic gradient step and noisy estimations of the natural gradient of the variational objective with respect to $\alpha_{kj}$. Following the computational method of the natural gradient (Hoffman et al. 2013), we compute the natural gradient of equation (1) with respect to the global variational parameters of $\alpha_{kj}$.
We consider Dirichlet distribution as a prior distribution with concentration parameters $\lambda_{kj}$, so a conditional distribution of profile given the observation data has form of the Dirichlet distribution. We consider the representation of the exponential family,

$$p(\alpha|Z,\phi,\gamma,\epsilon,\beta) = h(\alpha)\exp\left(\eta(Z,\phi,\gamma,\epsilon,\beta)^T t(\alpha) - a(\eta(Z,\phi,\gamma,\epsilon,\beta))\right)$$

where: $h(.)$ is the base measure; $a(.)$ is the log-normalize; $\eta(.)$ is the natural parameter; $t(.)$ is the sufficient statistics.

As above assumption of the variational parameters, we set $q(\alpha|\lambda^*)$ to be Dirichlet distribution as the complete conditional distributions. So we get

$$q(\alpha|\lambda^*) = h(\alpha)\exp\left((\lambda^*)^T t(\pi) - a(\lambda^*)\right)$$

We consider the lower bound for only $\alpha$,

$$\begin{aligned}\mathcal{L}(\alpha) &= \mathrm{E}_Q\left[\log p(\alpha|Z,\phi,\gamma,\epsilon,\beta)\right] - \mathrm{E}_Q\left[q(\alpha|\lambda^*)\right]\\ &= \mathrm{E}_Q\left[\log\{h(\alpha)\} + \eta(Z,\phi,\gamma,\epsilon,\beta)^T t(\alpha) - a(\eta(Z,\phi,\gamma,\epsilon,\beta)) - \log\{h(\alpha)\} - (\lambda^*)^T t(\alpha) + a(\lambda^*)\right]\\ &= \mathrm{E}_Q\left[\eta(Z,\phi,\gamma,\epsilon,\beta)^T t(\alpha) - a(\eta(Z,\phi,\gamma,\epsilon,\beta)) - (\lambda^*)^T t(\alpha) + a(\lambda^*)\right]\\ &= \mathrm{E}_Q\left[\eta(Z,\phi,\gamma,\epsilon,\beta)\right]^T\left[\nabla_{\lambda^*}\{a(\lambda^*)\}\right] - a(\eta(Z,\phi,\gamma,\epsilon,\beta)) - (\lambda^*)^T\left[\nabla_{\lambda^*}\{a(\lambda^*)\}\right] + a(\lambda^*)\end{aligned}$$

where, the expected value of the sufficient statistics is the gradient of log normalizer $\mathrm{E}_Q[t(\alpha)] = \nabla_{\lambda^*}\{a(\lambda^*)\}$
If the classical principal of the gradient method for maximization is used directly to find a maximum of $\mathcal{L}(\alpha)$ based on taking step of size $\rho$ in direction of the gradient, the optimized results is

$$\lambda_{(t+1)}^* = \lambda_{(t)}^* + \rho\nabla_{(\lambda^*)}L(\alpha) = \lambda_{(t)}^* + \rho^{(t)}\left[\nabla_{(\lambda^*)}^2\{a(\lambda^*)\}\right]\left\{E_Q\left[\eta(Z,\phi,\gamma,\epsilon,\beta)\right]^T - (\lambda^*)^T\right\}$$

Following (Hoffman et al. 2013), by premultiplying the gradient by the inverse Fisher information $\mathrm{G}(\lambda^*)$ and apply the stochastic natural gradient of the variational objective with respect to $\alpha$, we get

$$G(\lambda^*) = E_{\lambda^*}\left[\left(\nabla_{(\lambda^*)}\log q(\alpha|\lambda^*)\right)\left(\nabla_{(\lambda^*)}\log q(\alpha|\lambda^*)\right)^T\right] = \nabla_{(\lambda^*)}^2\{a(\lambda^*)\}$$

$$\widehat{\nabla_{(\lambda^*)}}L(\alpha) = \{G(\lambda^*)\}^{-1}\nabla_{(\lambda^*)}L(\alpha) = \{E_Q[\eta(Z,\phi,\gamma,\epsilon,\beta)] - \lambda^*\}$$

$$\begin{aligned}&\lambda^*_{(t+1)} = \lambda^*_{(t)} + \rho\widehat{\nabla_{(\lambda^*)}}L(\alpha) = \lambda^*_{(t)} + \rho\left\{E_Q\left[\eta\left(Z,\phi,\gamma,\epsilon,\beta\right)\right] - \lambda^*_{(t)}\right\}\\&= (1-\rho)\,\lambda^*_{(t)} + \rho\left\{E_Q\left[\eta\left(Z,\phi,\gamma,\epsilon,\beta\right)\right]\right\}\end{aligned} \tag{6}$$

Based traditional variational inference, we get the conditional distribution of $\alpha$

$$\log p\left(\alpha|Z,\phi,\gamma,\epsilon,\beta\right) = \sum_{i=1}^{N}\sum_{j=1}^{S}\sum_{k=1}^{K_{max}}\log\left(\frac{\Gamma(X_{ij}+\alpha_{kj})}{\Gamma(\alpha_{kj})}\right)\times\log\left(\frac{\Gamma(X_{ij}+\alpha_{kj})}{\Gamma(\alpha_{kj})}\right)\times \mathrm{E}_Q\left[Z_{ik}\right]\mathrm{E}_Q\left[\phi_{ij}\right] + \log\left(\alpha_{kj}\right)\left(\lambda_{kj}-1\right) + const$$

$$\mathrm{E}_q\left[\eta\left(Z,\phi,\gamma,\epsilon,\beta\right)\right] = \sum_{i=1}^{N}\mathrm{E}_Q\left[z_{ik}\right]\mathrm{E}_Q\left[\phi_{ij}\right]\overline{\alpha_{kj}}\left[\Psi\left(\sum_{j=1}^{S}\overline{\alpha_{kj}}\right) - \Psi\left(\sum_{j=1}^{S}X_{ij} + \sum_{j=1}^{S}\overline{\alpha_{kj}}\right) + \Psi\left(\overline{\alpha_{kj}} + X_{ij}\right) - \Psi\left(\overline{\alpha_{kj}}\right)\right] \tag{7}$$

By substituting equation (7) into equation (6), the final result of updated variational parameters is

$$\begin{aligned}&\left(\lambda^*_{kj}\right)^{(t+1)} = \left(1-\rho^{(t)}\right)\left(\lambda^*_{kj}\right)^{(t)}\\&+\rho^{(t)}\left\{\lambda_{kj} + \textstyle\sum_{i=1}^{N} r_{ik}f_{ij}\overline{\alpha_{kj}}\left[\Psi\left(\sum_{j=1}^{S}\overline{\alpha_{kj}}\right) - \Psi\left(\sum_{j=1}^{S}X_{ij} + \sum_{j=1}^{S}\overline{\alpha_{kj}}\right) + \Psi\left(\overline{\alpha_{kj}} + X_{ij}\right) - \Psi\left(\overline{\alpha_{kj}}\right)\right]\right\}\end{aligned}$$

#### 2.4 Updating variational parameters of $\beta$

Similarity, the stochastic natural gradient of the variational objective with respect to $\beta$

$$\begin{aligned}&\left(\iota^*_j\right)^{(t+1)} = \left(1-\rho^{(t)}\right)\left(\iota^*_j\right)^{(t)}\\&+\rho^{(t)}\left\{\iota_j + \textstyle\sum_{i=1}^{N}\left[1-f_{ij}\right]\overline{\beta_j}\left(\left[\Psi\left(\sum_{j=1}^{S}\overline{\beta_j}\right) - \Psi\left(\sum_{j=1}^{S}X_{ij} + \sum_{j=1}^{S}\overline{\beta_j}\right)\right] + \left[\Psi\left(\overline{\beta_j} + X_{ij}\right) - \Psi\left(\overline{\beta_j}\right)\right]\right)\right\}\end{aligned}$$

#### 2.5 Updating stick-breaking representation

Similarity, the optimized solution to the posterior distribution of unit length sticks $\gamma_k$

$$\begin{aligned}&\log Q^*\left(\gamma_k\right) = (1-1)\log\left(\gamma_k\right) + (\nu-1)\log\left(1-\gamma_k\right)\\&+\sum_{i=1}^{N}\mathrm{E}_Q\left[z_{ik}\right]\log\left(\gamma_k\right) + \sum_{i=1}^{N}\sum_{k'=k+1}^{K_{\max}}E_Q\left[z_{ik'}\right]\log\left(1-\gamma_k\right) + const\end{aligned} \tag{8}$$

which has the logarithmic form of the Beta distribution and the corresponding variational distributions of breaking proportions $q\left(\gamma_k\right)$ are considered to be the Beta distribution. Then, the standard conditions are satisfied for a closed form coordinate update for local parameters. Based on the stochastic natural gradient, the variational parameters of $\gamma_k$ are updated by computing variational expectations in equation (8) (Blei and Jordan. 2006, Hoffman et al. 2013),

$$\begin{aligned}&\left(\vartheta_k\right)^{(t+1)} = \left(1-\rho^{(t)}\right)\left(\vartheta_k\right)^{(t)} + \rho^{(t)}\left\{1 + \sum_{i=1}^{N}r_{ik}\right\}\\&\left(\vartheta'_k\right)^{(t+1)} = \left(1-\rho^{(t)}\right)\left(\vartheta'_k\right)^{(t)} + \rho^{(t)}\left\{\nu + \sum_{i=1}^{N}\sum_{k'=k+1}^{K_{\max}}r_{ik'}\right\}\end{aligned}$$

#### 2.6 Updating variational parameters of $\epsilon$

Similarity, the stochastic natural gradient of the variational objective with respect to $\epsilon$

$$\begin{aligned}&\left(\xi^*_{j1}\right)^{(t+1)} = \left(1-\rho^{(t)}\right)\left(\xi^*_{j1}\right)^{(t)} + \rho^{(t)}\left\{\xi_{j1} + \textstyle\sum_{i=1}^{N}f_{ij}\right\}\\&\left(\xi^*_{j2}\right)^{(t+1)} = \left(1-\rho^{(t)}\right)\left(\xi^*_{j2}\right)^{(t)} + \rho^{(t)}\left\{\xi_{j2} + \textstyle\sum_{i=1}^{N}\left(1-f_{ij}\right)\right\}\end{aligned}$$

#### 3 Reference


1. Bishop. C. M. 2006. Pattern Recognition and Machine Learning. Springer, 1 edition.

2. Blei, D and Jordan, M. 2006. Variational inference for Dirichlet process mixtures. Journal of Bayesian Analysis, 1:121–144.

3. Gopalan, P., Hao, W., Blei, D., Storey, J. 2016. Scaling probabilistic models of genetic variation to millions of humans. Nat Genet. 48, 1587-1590.

4. Hoffman, M., Blei, D., Wang, C. & Paisley, J. 2013. Stochastic variational inference. J. Mach. Learn. Res. 14, 1303–1347.

5. Jordan, M., Ghahramani, Z., Jaakkola, T. & Saul, L. 1999. Introduction to variational methods for graphical models. Mach. Learn. 37, 183–233.

6. Lartillot N. 2006. Conjugate Gibbs sampling for Bayesian phylogenetic models. J. Comput. Biol. 13:1701–1722.

7. Robbins, H. and Monro, S. 1951. A stochastic approximation method. The Annals of Mathematical Statistics 22, 400–407.

8. Wainwright, M. & Jordan, M. 2008. Graphical models, exponential families, and variational inference. Found. Trends Mach. Learn. 1, 1–305.
